# WGBS of Differentiating Adipocytes Reveals Variations in DMRs and Context-Dependent Gene Expression

**DOI:** 10.1101/2024.03.14.583264

**Authors:** Binduma Yadav, Dalwinder Singh, Shrikant Mantri, Vikas Rishi

**Author notes:** Corresponding authors:, Website: www.nabi.res.in. Contributed equally (Binduma Yadav, Dalwinder Singh).

## Abstract

Obesity, characterised by the accumulation of excess fat, is a complex condition resulting from the combination of genetic and epigenetic factors. Recent studies have found correspondence between DNA methylation and cell differentiation, suggesting a role of the former in cell fate determination. There is a lack of comprehensive understanding concerning the underpinnings of preadipocyte differentiation, specifically when cells are undergoing terminal differentiation (TD). To gain insight into dynamic genome-wide methylation, 3T3 L1 preadipocyte cells were differentiated by a hormone cocktail. The genomic DNA was isolated from undifferentiated cells and 4 hrs (4H), 2 days (2D) post-differentiated cells, and 15 days (15D) TD cells. We employed whole-genome bisulfite sequencing (WGBS) to ascertain global genomic DNA methylation alterations at single base resolution as preadipocyte cells differentiate. The genome-wide distribution of DNA methylation showed similar overall patterns in pre- and post- and terminally differentiated adipocytes, according to WGBS analysis. DNA methylation decreases at 4H after differentiation initiation, followed by methylation gain as cells approach TD. Studies revealed novel differentially methylated regions (DMRs) associated with adipogenesis. DMR analysis suggested that though DNA methylation is global, noticeable changes are observed at specific sites known as ‘hotspots.’ Hotspots are genomic regions rich in transcription factor (TF) binding sites and exhibit methylation-dependent TF binding. Subsequent analysis indicated hotspots as part of DMRs. The gene expression profile of key adipogenic genes in differentiating adipocytes is context-dependent, as we found a direct and inverse relationship between promoter DNA methylation and gene expression.

## Introduction

DNA methylation is a fundamental epigenetic mechanism that plays a vital role in cell differentiation, a process by which an undifferentiated cell becomes a more specialized cell type with specific functions and characteristics[1][2], [3][4]. DNA methylation is a chemical modification that involves adding a methyl group (CH_3_) to the cytosine base of CpG dinucleotides, where guanine succeeds cytosine[4][5][6][7]. This modification typically occurs at the 5’ carbon of the cytosine ring, and it is performed by DNA methyltransferases (DNMTs) [8]. CpG sites are found throughout the genome, and their methylation status can be heritable, allowing the epigenetic information to be transmitted from one cell generation to the following [9]. The consequences of its dysregulation further underscore the importance of DNA methylation in cell differentiation. Abnormal DNA methylation patterns have been implicated in various developmental disorders and diseases, including cancer[7][10][11]. Hypomethylation, the loss of DNA methylation, can lead to the reactivation of silenced genes, potentially causing cells to revert to an undifferentiated state or exhibit uncontrolled growth. Alternatively, hypermethylation, the excessive methylation of CpG sites, can result in the inappropriate silencing of critical genes, disrupting normal cellular differentiation processes[10][11][12][13][14].

A molecular mechanism is proposed on how DNA methylation in gene promoter regions can act as an active or repressive mark, allowing or inhibiting the binding of transcription factors and other regulatory proteins necessary for gene activation[15][16][17]. Conversely, DNA methylation in gene body regions is generally associated with gene activation. This process, known as gene body methylation, is less understood but appears to play a role in enhancing transcriptional elongation and stabilising gene expression levels[18][19].

DNA methylation plays a remarkable role in adipogenesis as in many physiological processes, the process in which preadipocytes (undifferentiated cells) develop into mature adipocytes (fat cells). During adipogenesis, multipotent mesenchymal stem cells (MSCs) differentiate into mature adipocytes through tightly regulated molecular events. The orchestration of this process involves epigenetic alterations such as DNA methylation, modifications in histones, and the regulation by non-coding RNA [20]. Previous studies have shown that dynamic alterations in DNA methylation patterns occur during adipogenesis, affecting the expression of genes associated with adipocyte development, lipid metabolism, and adipose tissue function. For example, changes in methylation status at promoters of adipogenic transcription factors (e.g., PPARγ and C/EBPα) can influence their expression levels, which in turn drive the expression of adipocyte-specific genes, such as that encoding adiponectin, leptin, and fatty acid-binding protein 4 (FABP4), thereby impacting adipocyte differentiation[21][22][23][24][25][26]. The role of DNA methylation in adipogenesis can be observed at three stages of the process: 1) preadipocyte commitment, 2) Early differentiation, and 3) Late differentiation. However, how DNA methylation selectively regulates and changes its pattern during adipogenesis, thus leading to changes in gene expression, needs to be studied in detail[27]. Because the 3T3-L1 cell line exhibits a distinct and synchronised differentiation process from pre-adipocytes to fully grown, lipid-laden adipocytes that mimic preadipocyte differentiation *in vivo*, it is considered an appropriate model for studying adipogenesis. Environmental cues like diet and hormonal treatment initiate differentiation, causing or leading to significant chromatin remodelling and epigenomic changes, beginning 4H after induction and proceeding to TD[28][29][30]. Also, two distinct waves of transcription factors starts off adipogenesis[28]. We used WGBS with one base resolution to demonstrate how genome-wide methylation patterns changed as 3T3 L1 cells differentiated by hormonal treatment. We focused on four essential time points: Pre-AD; day 0), representing the initial period when adipogenic factors are relatively inactive; 4H after differentiation induction, when the first wave transcription factors are highly active while the second wave transcription factors are expressed low; 2D, which marks the beginning of the elevated expression of the second wave of transcription factors and the initiation of terminal differentiation, and 15D, which displays the fully developed, mature, and lipid-rich adipocyte[31]. To our current understanding, this is the first attempt to analyse genome-wide methylation patterns in TD cells. Furthermore, we have looked for site-specific methylation at transcription factor hotspots where multiple transcription factors bind cooperatively and modify the structure of chromatin within hours after the induction of adipogenesis. Studying the epigenetic profile of preadipocytes and adipocytes can contribute in developing various treatments for obesity and metabolic disorders such as treating genetic disorders and other metabolic disorders through *ex vivo* gene therapy utilizing preadipocytes[32]. Unrevealing the mechanisms by which DNA methylation regulates differentiation will improve our understanding of the underlying biological pathways.

## Materials and methods

### Chemicals and reagents

Dulbecco’s Modified Eagle’s Medium (DMEM) and fetal bovine serum (FBS) were purchased from Gibco, lnc. (Grand Island, NY). Isobutylmethylxanthine (IBMX), dexamethasone, insulin, and 3-(4, 5-dimethylthiazol-2-yl)-2, 5-diphenyl tetrazolium bromide (MTT) were obtained from Sigma (St. Louis, MO). The EZ DNA Methylation Kit from Zymo Research was used for the bisulfite treatment of DNA (Zymo Research, Cat. No. D5001, USA).

### 3T3-L1 Preadipocytes culture and differentiation

3T3-L1 preadipocytes obtained from NCCS, Pune, India, were cultured in DMEM and supplemented with 10% FBS. The cells were maintained at 37°C in a humidified atmosphere with 5% CO_2_. The cells were seeded at a density of 1×10^5^ cells/ml in a 6-well culture plate. After two days of reaching confluence, cell differentiation was induced by treating the cells with a differentiation medium containing 10% FBS DMEM supplemented with an MDI hormone cocktail (0.5μM isobutylmethylxanthine IBMX, 5μM dexamethasone, and 0.5μg/ml insulin). The medium was then replaced with 10% FBS DMEM containing 5 μg/ml insulin. Finally, the differentiation medium was replaced with 10% FBS DMEM. The 3T3-L1 preadipocytes were divided into four groups based on the post-induction time: undifferentiated preadipocytes, 4H and 2D post-induction, and 15D post-induction when cells are considered fully or terminally differentiated.

### Genomic DNA isolation

The cells were harvested and washed twice with cold PBS. Subsequently, the cell pellets were snap-frozen in liquid nitrogen for storage at -80°C or further processed. For DNA extraction, the frozen cell pellets were thawed at room temperature and resuspended in PBS following the instructions provided by the manufacturer (DNeasy Blood & Tissue Kits, Cat. No. 69504). Extracted DNA was further checked for quality on gel and subsequently bisulfite-treated and was used for WGBS sequencing and cloning of CpG-rich regions and DMRs.

### PCR amplification, hotspot cloning, and bisulfite treatment of samples for sequencing

Hotspots were cloned to examine their DNA methylation states as the cells differentiate. Purified genomic DNA from undifferentiated and differentiated adipocytes was subjected to bisulfite modification using the EZ DNA Methylation kit (Zymo Research). Approximately 0.5-1μg of genomic DNA was treated with bisulfite and eluted in 20μl elution buffer following the manufacturer’s protocol. After bisulfite treatment, 2μl of the eluted DNA was amplified for 40 cycles using methylation-specific primers according to standard protocols. The PCR products were visualized by agarose gel electrophoresis and extracted from the gel using a gel extraction kit (Qiagen, Cat. No. 286040). The PCR products obtained from the gel were then cloned into the pcDNA plasmid as BamHI-XhoI fragments. Plasmid DNA was isolated from individual clones using the QIAprep Spin miniprep kit (Qiagen, Cat. No. 27106), and the cloned plasmids were subjected to sequencing using T7 forward primer and SP6 reverse primer designed for the vector backbone. The sequencing data provided information on cytosine methylation at each CpG site within the amplicon. The chromatograms obtained from sequencing were analyzed using Snapgene software, and the sequencing data were further analyzed for DNA methylation using BIQ Analyser software.

### Library Preparation for WGBS

High-quality genomic DNA was extracted using standard phenol/chloroform extraction, ethanol precipitation, or the DNeasy Blood and Tissue kit. The preadipocytes and adipocytes were lysed in lysis buffer at 37°C for 1hr and then digested with proteinase K at 10μg/ml concentration for 3hrs at 50°C. Following cell lysis, DNA was isolated using the phenol-chloroform extraction method. Eurofins Genomic India Pvt. Ltd. provided bisulfite conversion and sequencing services. To confirm the efficiency of bisulfite conversion, lambda DNA spike-in was added, and it was found that 99% of the DNA was successfully bisulfite converted. For library construction, 100ng of genomic DNA was treated with the EZ DNA Methylation-Gold kit (Zymo Research) for bisulfite conversion. The resulting libraries, consisting of DNA fragments with lengths between 200-400 bps, were subjected to 150 bps pair-end sequencing on an Illumina platform. All sequencing analyses were performed based on the *Mus musculus* NCBI GRC38 genome assembly (mm10 version). The sequencing statistics can be found in FigureS1. The raw WGBS data (FASTQ and bedGraph files) is deposited in the NCBI SRA database.

### RNA Isolation and Quantitative real-time RT-PCR (RT-qPCR)

Total RNA was extracted from 3T3-L1 preadipocytes, 4H post-differentiation, 2D post-differentiation, and 15D TD cells by <u>TRIzol</u> reagent (Ambion, USA). The purity and concentration of isolated RNAs were determined using the NanoDrop spectrophotometer (Thermo Fisher Scientific, USA). cDNA was synthesised by the iScript^TM^ cDNA synthesis kit (Bio-Rad Laboratories, Inc). The mRNA expression of 45 adipogenic genes, i.e., KLF5, KLF6, STAT5a, ZFP423, ZFP467, Tcf7l1, KLF2, Foxo1, Foxa2, Foxc2, Cd36, Lpl, Fasn, Plin1, Plin2, Plin3, Plin4, Plin5, DGAT1, ANGPTL4, PDGFRα, PDGFRβ, VEGFc, VEGFb, EGR2, Resistin, Lipoproteinlipase, FABP4, CREB1, TET1, TET2, TET3, ADIPOQ, GATA2, EBF1, HOXA6, HOXA5, VDR, KLF4, ATF7, JUNB, PBX1, Slc2a1/GLUT1, KLF14 was evaluated using real-time PCR on CFX96 Real-Time system with SYBR Green Fast qRT-PCR mix from Bio-Rad. The reaction protocol involved priming at 25°C for 5min, reverse transcription at 46°C for 20 min, and RT inactivation at 95°C for 1min. The gene expression levels were calculated using the normalised relative quantification protocol followed by the 2^-ΔΔCT^ method.

### WGBS Data analysis

The quality of raw sequences was examined with FastQC (v0.11.9), and Trimmomatic (v0.39) [33] was used to remove the Illumina adaptor sequences and to filter out the low-quality reads and bases (Phred quality score < 15) using “SLIDINGWINDOW:4:15 LEADING:3 TRAILING:3 MINLEN:36 HEADCROP:10 ILLUMINACLIP: TruSeq3-PE.fa:2:30:10” parameters (Table 1). Following, clean reads were mapped to the mm10 (GRCm38) reference genome using Bismark (v0.23.1)[34]. The lambda genome (GenBank: J02459.1) was also mapped along with the reference genome to determine the bisulfite conversion efficiency. (Table 1 & FigureS1). The obtained bisulfite conversion rate for CG context was above 99% for all libraries (all samples)

**Table 1:** The paired-end mapping with Bismark was performed with Bowtie 2 using the following parameters: –score_min L, 0, -0.6 -X 1000, and duplicated reads are removed using the deduplicate_bismark command (Figure S1C). The genome-wide cytosine analysis was performed using the remaining reads; its results are given in (Figure S1D). The methylation bias in the reads was determined with the -mbias option of Bismark Methylation Extractor; consequently, methylated CpGs were extracted by ignoring one nucleotide of 3’end of both reads along with -no-overlap -comprehensive -bedGraph -cytosine_report options.

| Sample<br>s | Raw reads | Clean read | Clea<br>n<br>base<br>s (G) | Clea<br>n<br>ratio<br>(%) | Mapped<br>reads | Mapping<br>rate<br>(%) | Duplicatio<br>n rate (%) | Bisulphite<br>conversion<br>rate (%) |
| --- | --- | --- | --- | --- | --- | --- | --- | --- |
| Pre-AD | 55,438,326 | 53,411,178 | 14.5 | 87.2 | 39,346,076 | 73.7 | 19.9 | 99.3 |
| 4H | 51,941,175 | 42,412,105 | 11.6 | 74.6 | 31,456,049 | 74.2 | 18.1 | 99.3 |
| 2D | 55,463,283 | 53,311,291 | 14.4 | 86.7 | 45,153,417 | 84.7 | 18.5 | 99.4 |
| 15D | 52,949,471 | 49,723,797 | 13.4 | 84.2 | 36,544,292 | 73.5 | 23.5 | 99.2 |

**Table 1:** The paired-end mapping with Bismark was performed with Bowtie 2 using the 1109 following parameters: `-score_min L, 0, -0.6 -X 1000`, and duplicated reads are removed using 1110 the `deduplicate_bismark` command (Figure S1C).
| Sample<br>s | Raw reads | Clean<br>read | Cle<br>an<br>bas<br>es<br>(G) | Clean<br>ratio<br>(%) | Mapped<br>reads | Mapping<br>rate<br>(%) | Duplicatio<br>n rate (%) | Bisulphite<br>conversio<br>n rate (%) |
| --- | --- | --- | --- | --- | --- | --- | --- | --- |
| Pre-AD | 55,438,326 | 53,411,178 | 14.5 | 87.2 | 39,346,076 | 73.7 | 19.9 | 99.3 |
| 4H | 51,941,175 | 42,412,105 | 11.6 | 74.6 | 31,456,049 | 74.2 | 18.1 | 99.3 |
| 2D | 55,463,283 | 53,311,291 | 14.4 | 86.7 | 45,153,417 | 84.7 | 18.5 | 99.4 |
| 15D | 52,949,471 | 49,723,797 | 13.4 | 84.2 | 36,544,292 | 73.5 | 23.5 | 99.2 |

The Pearson’s correlation coefficient of pre-adipocyte and remaining samples was obtained using the MethylKit R package (v1.20.0)[35]. DMRs between control (pre-adipocytes) and 4H, 2D, and 15D were detected using the DSS package (2.42.0)[36]. In the pairwise comparison of control versus rest, DMLtest function with a smoothing span of 100 bps was applied to estimate mean methylation levels, and the callDMR function was used to detect DMRs having minimum 3 CpG sites, methylation difference >20%, minimum 50 bp length, and p-value<0.05. Further, the DMRs, which are 100 bp apart, are merged.

For downstream analysis, the obtained DMRs were categorised into hypo- and hyper-methylated (GO-Gene Ontology and KEGG-Kyoto encyclopedia of genes and genomes analysis). The annotation was performed with ChIPseeker R package (v1.30.3)[37] and RefSeq mm10 annotation (http://hgdownload.cse.ucsc.edu/goldenpath/mm10/bigZips/genes/). The annotatePeak function of ChIPseeker was used to annotate hypo- and hyper DMRs by defining the promoter as 3kb upstream of the transcription start site (TSS). The obtained annotated genomic features, such as promoter, UTRs, exons, introns, and intergenic regions, were used for comparison and visualisation. GO enrichment analysis of genes whose promoter overlapped with DMRs was performed by R package clusterProfiler (v4.2.2) with enrichGO function. The GO terms were determined based on the default Benjamini-Hochberg (BH) procedure and a cutoff score of adjusted p-value <0.01 or q-value<0.05, depending on the selected parameters. Further, enrichKEGG of clusterProfiler function is used for pathway analysis with the BH procedure (final parametric values were produced with fixed p-value cutoff = 1, p-AdjustMethod = “BH”, minGSSize = 1, maxGSSize = 500, q-value cutoff = 1).

### Genomic region analysis

The RefSeq genes annotation of mouse reference genome mm10 (GRCm38) was obtained from UCSC (https://hgdownload.soe.ucsc.edu/goldenPath/mm10/bigZips/genes/), and the mouse genome was divided into 9 regions. To avoid redundancy for protein-coding genes with multiple transcripts, only the longest was used for defining the locations of promoters, TSS, TES, exons, introns, and intergenic regions[38][39]. Promoters are described at 0-3000 bases upstream of the TSS, 5’ untranslated region (UTR) between the TSS and ATG start site, gene body between ATG and stop codon, all exons in the gene body, first exon of gene body, all introns in the gene body, first intron in the gene body, 3’UTR between the stop codon and poly-A site (or end of TSS), and intergenic regions as remaining regions between two genes[40]. Additionally, genomic locations of CpG islands and RepeatMasker were downloaded from the ‘UCSC table browser (https://genome.ucsc.edu/cgi-bin/hgTables). The regions associated with CpG islands (CGI) were also explored by considering both shores (0-3000bp in the upstream and downstream of CGI) and shelves (3000-4000bp) in the upstream and downstream of CGIs.

DeepTools suite (3.5.1)[41] was used to generate and plot average methylation levels of different genome regions. The computeMatrix scale-regions were used to measure mean methylation levels across non-overlapping windows with the following parameters: --binSize 10 --numberOfProcessors 40 --regionBodyLength 3000 -b 2000 -a 2000 for promoters and -b 0 -a 0 for other genomic elements or features. The plotProfile was used to compute the data matrix required for visualisation. The bedGraph files obtained from the Bismark methylation extraction step were converted into bigwig format using UCSC bedGraphToBigWig for processing in DeepTools. Circos and Gene chromosome plots were made using LaTex with in-house scripts.

### Statistical analysis

Data was analysed using Excel and GraphPad Prism and presented as mean ± SEM. P < 0.05 was considered significant.

## RESULTS

### DNA methylation profile during preadipocyte differentiation

Data analysis revealed that DNA methylation exhibits changes throughout the adipocyte cell lineage, occurring during and after the differentiation process.

To investigate the whole-genome DNA methylome profiles associated with lineage-specific adipogenesis, 3T3-L1 cells were cultured and induced to differentiate from preadipocytes to mature adipocytes *in vitro*. We performed WGBS on 3T3-L1 preadipocytes (Pre-AD) and differentiated cells by extracting and analysing genomic DNA at 4H, 2D, and 15D post-differentiation (Figure S2). Images of undifferentiated and differentiated adipocytes are shown (Figure 1A).

**Figure 1:**
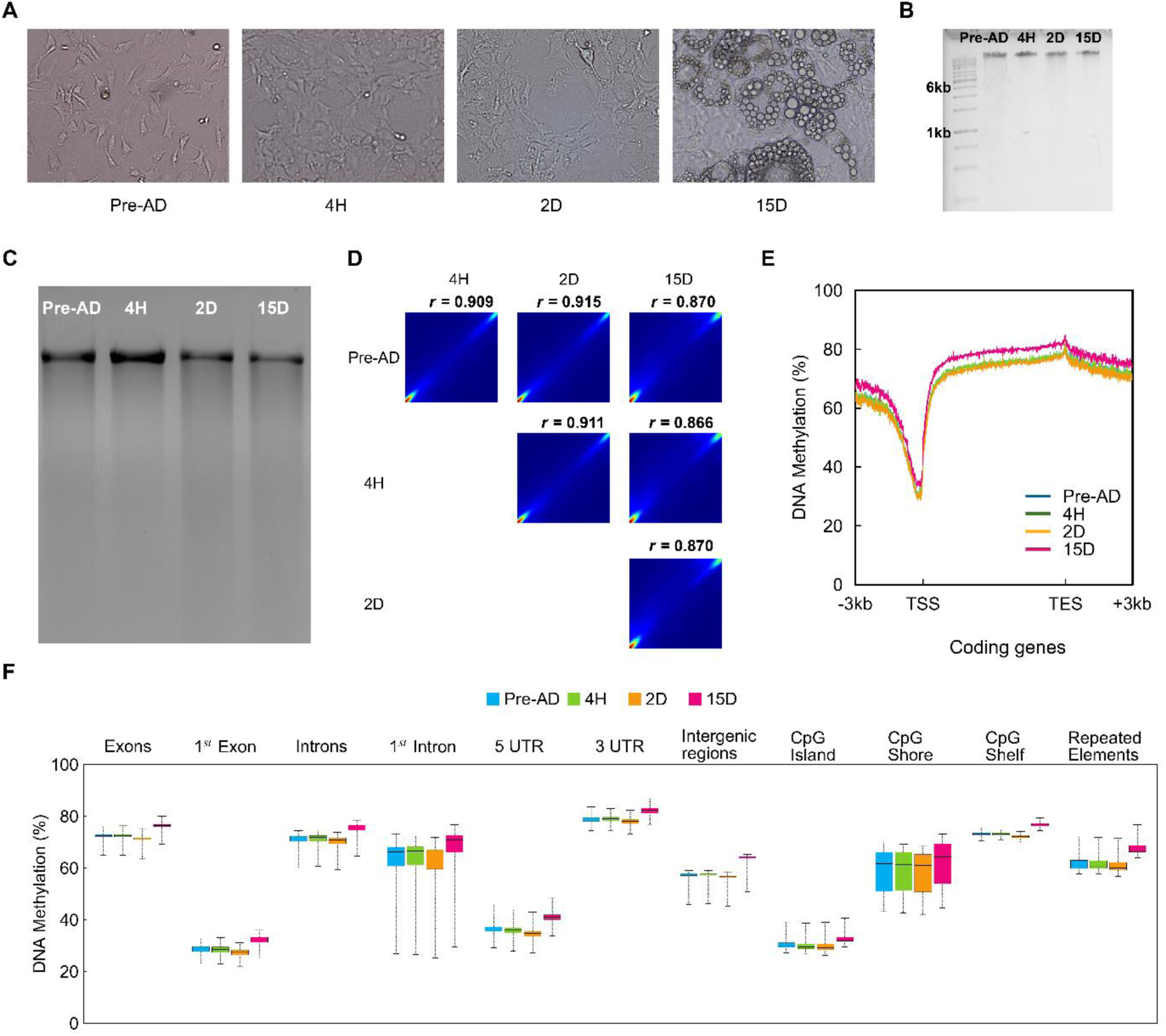
DNA methylation pattern during differentiation of preadipocytes to mature adipocytes. (A) Bright-field images depicting control 3T3-L1 preadipocytes and cells exposed to hormone cocktail to induce differentiation. Panels show Pre-AD (Preadipocytes), 4H (4hrs), and 2D (2 days) post-induction and 15D (15 days) post-induction and terminally differentiated (TD) cells. The presence of conspicuous lipid droplets characterises TD cells. (B) Genomic DNA extracted from 3T3-L1 cells shows genome integrity during the differentiation process. (C) McrBc restriction digestion of genomic DNA extracted from preadipocytes and 4H, 2D, and 15D post-induction suggest genome-wide loss and gain of DNA methylation. (D) Pearson’s correlation coefficient analysis of DNA methylation between preadipocytes and differentiating adipocytes and TD cells. (E) The DNA methylation levels with reference to the Transcription Start Site (TSS) and Transcription End Site (TES) in coding transcripts. Traces for the control sample are superimposed by 4H and 2D sample traces and are not shown. (F) The levels of DNA methylation at various genomic annotations such as exon, intron, 5’UTR, 3’UTR, CpG islands, CpG shores, and CpG shelves. RefSeq mm10 annotations were used to obtain transcripts. Promoters are defined by considering 3kb upstream regions. The bin size is 5, and the minimum base level depth of CpGs is 1.

Extracted genomic DNA samples were digested with the McrBc restriction enzyme, which cleaves methyl CpG-rich DNA (one or both strands) to ascertain the global DNA methylation in undifferentiated and differentiated cells (Figure 1B-C)[9]. At 4H, the genomic DNA band is intense compared to the faint band of undifferentiated and post-differentiated 2D and 15D samples. This observation is interpreted as depicting the hypomethylation of the genome at 4H post-differentiation. Examining the experiment’s fidelity and the sample selection’s rationality is crucial, and one key indicator is the correlation of methylation levels across samples. We conducted Pearson’s correlation coefficient analysis among samples, focussing on CG contents. The Pearson correlations between samples vary from R^2^ = 0.86-0.91, suggesting a strong correlation and lack of any substantial changes in DNA methylation among different samples (Figure 1D) except TD cells in which methylation is more pronounced.

WGBS analysis indicated that the overall global DNA methylation was similar inbetween preadipocytes and adipocytes (Figure 1E). Within the vicinity of the transcription start sites, a valley depicting the loss in methylation was observed in all four samples (Figure 1E). In contrast, higher DNA methylation was observed in gene body regions (Figure 1F), a common feature observed in various cell types[18][19][42]. We compared in-house (NABI dataset) WGBS data analysis with the adipogenic reprogramming dataset[43] to validate our WGBS data further (supplementary data depicting comparative analysis and correlation analysis between AR and NABI datasets Table S3, Figure S6, Figure S7, Figure S8, Table S4). The genome-wide distribution of DNA methylation exhibited similar patterns before and after the differentiation of adipocytes[44]. Nevertheless, when specifically considering methylated CpGs, a decline in trend was noted during adipocyte differentiation at the initial stages, suggesting DNA methylation is reduced in a restricted number of specific regions, indicating a regulatory role [45].

Furthermore, when whole genome regions were categorised as per different genomic annotations, preadipocytes and adipocytes exhibited variable DNA methylation patterns (Figure 1F). Also, the methylation pattern in all the chromosomes was congruent with the methylation levels in the coding regions (Figure S3). The intergenic regions showed a substantial increase in the DNA methylation level at 15D (Figure 1F). The process of terminal differentiation involves activating specific transcription factors and epigenetic modifications that regulate gene transcription, leading to the establishment of distinct cell fates[14]. To investigate the active demethylation process in 3T3-L1 preadipocytes, we compared the methylation levels between undifferentiated cells and differentiated cells. We classified a CpG site as demethylated if its methylation level decreased by more than 0.1 between the two compared stages, with statistical significance determined by Fisher’s exact test (p-value < 0.05; FDR < 10%). These results indicate active demethylation at many CpG sites during the transition from preadipocytes to mature adipocytes.

Our analyses revealed that some specific CpGs in preadipocytes are methylated after differentiation. Furthermore, a significant portion of highly methylated CpGs was found in introns, repeat regions, and gene bodies (Figure 1E). In contrast, most unmethylated CpGs were located in promoters and CpG islands (CGIs) (Figure 1E, F), suggesting the importance of maintaining these regions in an unmethylated state for gene expression. We also demonstrated that non-CpG cytosine methylation is also dynamic during preadipocyte differentiation (Figure S1D). Figure 1F illustrates the average methylation levels of various functional genomic elements in preadipocytes and adipocytes. Such an analysis revealed significant demethylation in several functional elements, including CGIs and 5’ untranslated regions (5’-UTRs). A similar DNA methylation pattern was observed in the chromosome-wise plot (Figure S3). Interestingly, it was further observed that the methylation status of CGIs near the TSS remained stable. In contrast, CGIs within genic regions displayed greater dynamism during the early stages of differentiation [46]–[48]. Our study offers insights into the DNA methylation patterns associated with lineage-specific adipogenesis, highlighting the dynamic nature of DNA methylation and its potential role in regulating gene expression during adipocyte differentiation.

### Site-specific methylation pattern at hotspots

Genome-wide DNA methylation pattern demonstrated variability. To check DNA methylation at specific locations that acted as hotspots, i.e., have sites for multiple transcription factors were evaluated for varying degrees of methylation at different time points used in this study. For example, CpG-rich hotspots on chromosome 5 and chromosome 8 were PCR amplified and bisulfite treated, cloned, and sequenced (Figure 2A, B). It further demonstrated that as the cells differentiated, the methylation pattern changed in the hotspot regions containing binding sites for adipogenic transcription factors[49]. This confirmed that cytosine methylation at four indicative periods was dynamic at the genome and site-specific level, emphasising the importance of cis-elements DNA methylation in gene regulation[50][51].

**Figure 2:**
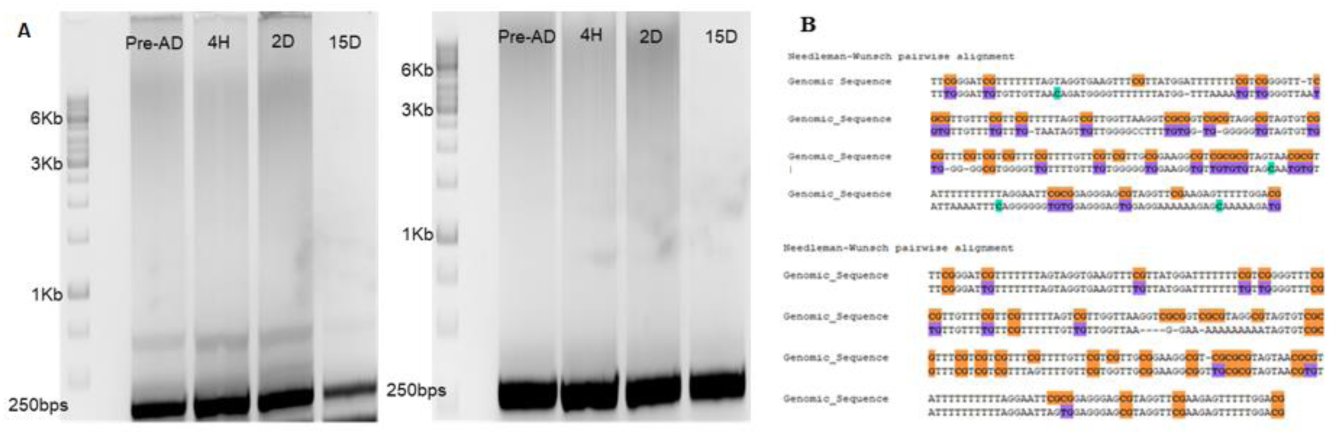
PCR amplification of hotspots at chromosomes 5 (chr5) and 8 (chr8). Methylation-independent primers were designed for the DNA methylation status of hotspots (250 bps) present at chr8: 19784539:19784789 and chr5: 13993644:139936704 using the Bisearch tool. (A) PCR amplified hotspots were bisulfite treated, then cloned in pcDNA plasmid, and Sanger sequenced. (B) Sanger sequence analysis of Bisulphite treated cloned hotspot with BIQanalyzer depicting the change in methylation pattern at specific sites.

### Characterization of DMRs

DMRs are contiguous genomic regions-with variable DNA methylation levels that differ between phenotypes[52][53]. DMRs can be found across the entire genome, but they are specifically recognized in and around gene promoter regions, within the gene bodies, and at intergenic regulatory regions[54][55][56][2][57][58][59][60]. These genomic regions are considered potential functional regions involved in the transcriptional control of genes since they exhibit varying levels of methylation across various samples (tissues, cells, etc.)[61]. Finding DMRs across several tissues may reflect the epigenetic basis of gene regulation between tissues and cells [45]. Numerous DMRs have been identified during developmental reprogramming stages[62]. Here, DMRs between control (pre-adipocytes) and 4H, 2D, and 15D were identified using the DSS package (2.42.0).

By analyzing a subset of known and unknown DMRs in 3T3-L1 cells, we identified a small proportion that exhibited changes in DNA methylation during differentiation (Figure 3). DMRs were categorised into hypo- and hyper-DMRs based on varying DNA methylation[63][62][64]. Comparing undifferentiated 3T3-L1 cells with 4H, 2D, and 15D fully differentiated cells, we found 200, 492, and 1036 novel DMRs, respectively. Furthermore, hypo-DMRs were dominant compared to hyper-DMRs at 4H and 2D post-differentiation, i.e., 51% and 56.9% (Figure 3A). At 15 days post-differentiation, the hyper-DMRs were predominant (88%) compared to hypo-DMRs (Figure 3A). We also compared the DNA methylation fold change in DMRs control vs. 4H, 2D, and 15D (Figure 3B-D). The circos plot demonstrated that at 4H post-differentiation, there was a pronounced hypomethylation (negative fold change), which suggests a significant reduction in methylation upon differentiation initiation. However, at 2D, there is substantial positive fold change depicting regain of DNA methylation. Furthermore, DMRs’ status varies. The significant fold change in hypo-DMRs was observed at chromosome 15 at 4H (Figure 3C), whereas significant changes were observed at chromosomes 2, 6, 5, 8, 11, 15, and 17 at 2D (Figure 3D) on chromosomes 5, 10, and 15. However, after 15D, the methylation level increased, indicating that hyper-DMRs are not uniformly distributed across all chromosomes. To summarise, the DMR distribution, whether hypo or hyper, is not uniform and is biased for a few chromosomes (Figure S5). Also, genomic annotations show that a significant fraction of the hyper-DMRs were enriched in distal intergenic regions (Figure 3E). The majority of DMRs were found in intronic and intergenic regions. In promoter regions, most DMRs were hypomethylated after 4H and hypermethylated after 15D.

**Figure 3:**
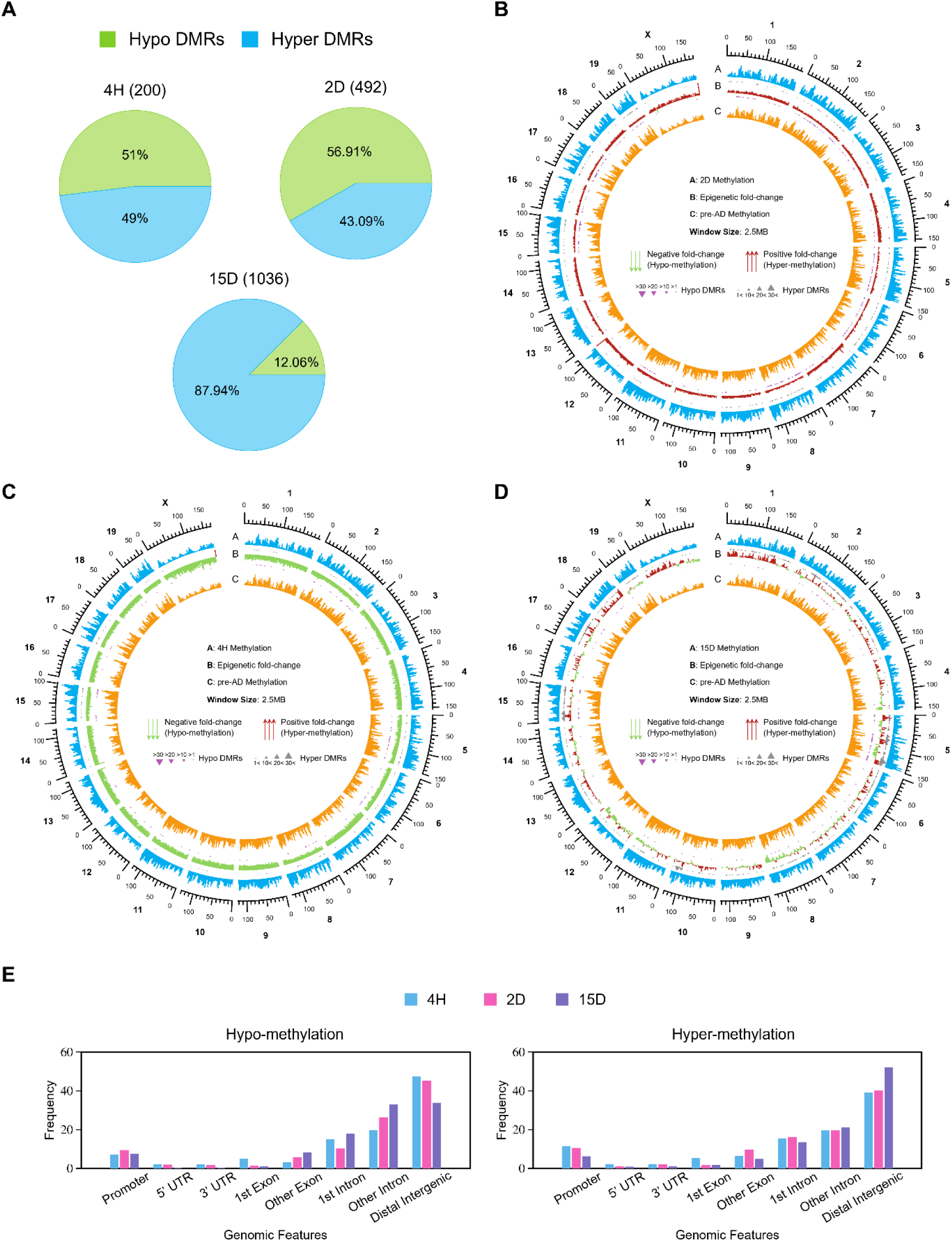
Analysis of DMRs present at different time points post-differentiation compared to undifferentiated cells. A) Pie chart depicting the difference between methylation level in hypo and hyper-DMRs at the three different time points: 4H, 2D, and 15D compared to control. B-D) Circos plot depicting fold change analysis of DMRs (B) control vs. 2D (C) control vs. 4H (D) control vs. 15D. E) Percentage of Hypo-DMRs and Hyper-DMRs at different time points post-differentiation at various genomic annotations. Windows: 5MB Overlapping: 2.5M Base level Depth:1

The study implies that the methylation alterations observed during differentiation were unidirectional, transient and involved both hypermethylation and hypomethylation. Also, prominent hypomethylation was seen at the induction of differentiation. These findings imply a complicated and dynamic underlying process involved in adipogenesis that depends on CpG methylation. Also, DMRs have an active role in gene regulation.

### Correlation between Hotspots and DMRs

We further overlapped the locations of hotspots and DMRs to understand the role of DMRs in gene regulatory activities. DMRs-containing binding motifs of major transcription factors that are part of hypoDMRs and hyperDMRs were investigated. To characterise DMRs of undifferentiated and differentiated adipocytes in an unbiased manner, the regions were subdivided into various genomic annotations such as 5’-UTR, 3’-UTR, CpG island, CpG shore, exons, and introns (Figure 3E). Few of the DMRs were part of hotspots, which suggests the regulatory role of DMRs during preadipocyte differentiation (Figure 4). A total of 200 (98 - hyper-DMR;102 -hypo-DMR), 492 (212 -hyper-DMR; 280 -hypo-DMR), and 1036 DMRs (911 -hyper-DMRs;125 -hypo-DMRs) were found by WGBS at 4H, 2D, and 15D, respectively and are found to overlap with 11974 hotspots (Figure 4A, B).

**Figure 4:**
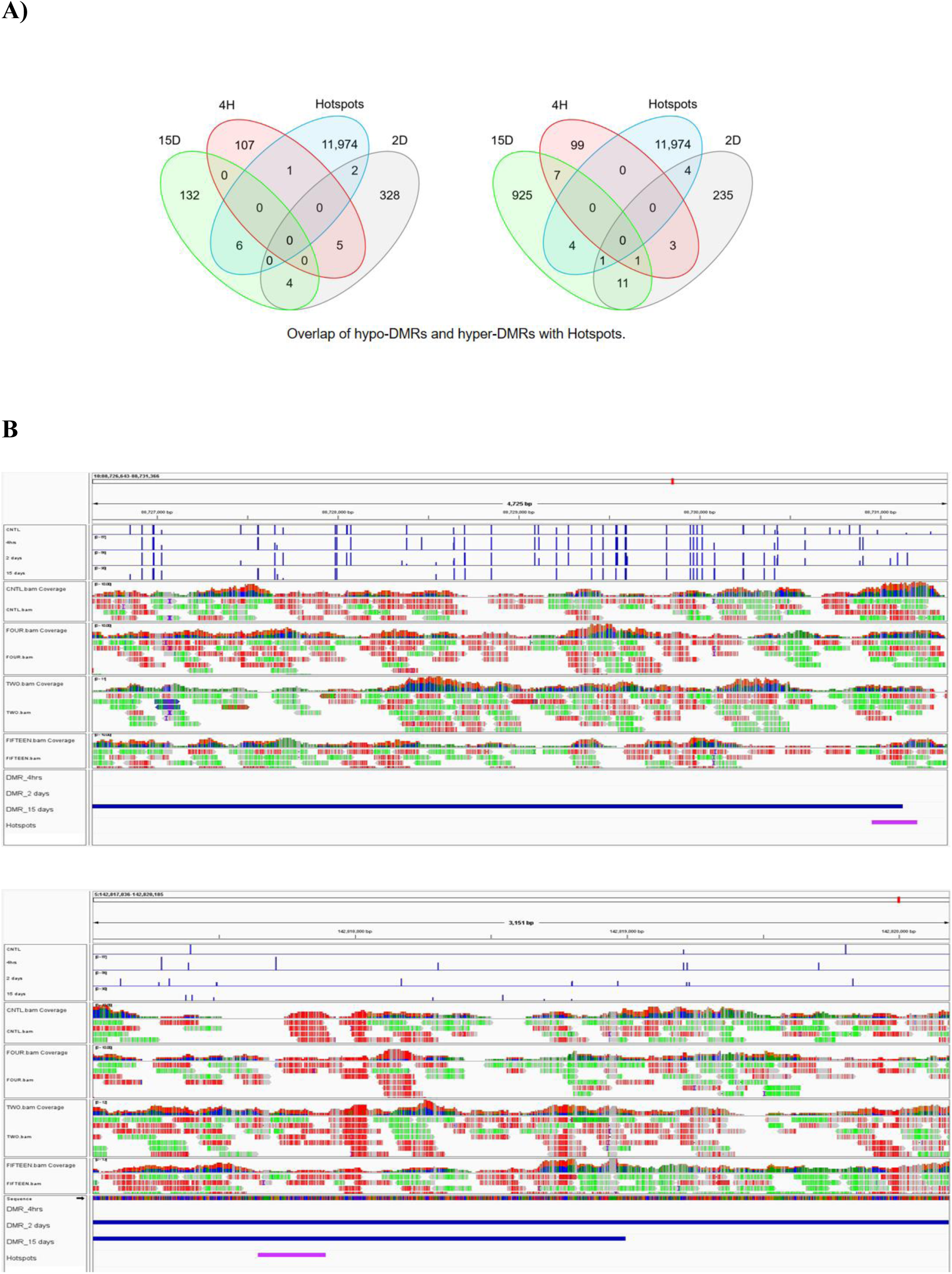
Hotspots are part of DMR. A) Venn diagram showing overlap patterns of DMRs and hotspots at indicated time points of differentiation. B) Representative genome browser view of overlap of Hotspots and DMRs using IGV (integrative genome viewer), which depicts Hotspots as a part of DMRs. Blue tracks represent DMRs at different time points, and pink track represents hotspots.

### DNA methylation variability in promoters

To investigate the probable mechanism for selective up or down-regulation of DNA methylation at loci unique to adipocytes, an integrative genome viewer (IGV) was used to evaluate the promoter DNA sequences of adipogenic genes showing complete coverage during 15D of terminal differentiation; there was an upturn in the average DNA methylation at the promoters of the genes MBD1, MBD3, DNMT3A, Lyz2, CD36, Tet1, GM10354, and GM14325(Figure5). In previous studies and based on qRT-PCR expression analysis here, entrenched upregulation of DNA methylation often leads to gene repression. However, in the case of Lyz2, upregulation in DNA methylation led to increased gene expression, suggesting a direct relationship or positive correlation exits between DNA methylation and gene expression. Lyz2 is responsible for lysozyme expression in 3T3-L1 cells, which maintains the expression of genes related to adipogenesis and adipocyte differentiation[65]. Similarly, as reported earlier, we observed elevated gene expression and elevated methylation in the VDR and EBF1 promoter regions on day 15D cells[43].

### Identification and Validation of differentially expressed genes

To identify differentially expressed genes, primers (Table S1) were designed using primer designing tools[66]. □-actin was considered an internal control to normalize gene expression signals (Table S1). Three biological replicates and two technical replicates for each stage were run along with internal control in qRT-PCR. Standard manufacturer protocol was used for the qRT-PCR reactions. The comparative cycle threshold (CT) method was used to determine the fold difference of the studied genes at each differentiation stage[67](Figure 6 and S4). One-way ANOVA was used to determine statistical differences between maturation stages (*p < 0.05). The adipogenic genes *ZFP467, Tet3, KLF4, CD36,* and *GM10354* were expressed 2 to 5 fold at initial differentiation. However, *Stat5a, VDR, EBF1, Lyz2,* and *Pbx174* showed an exponential increase in expression at terminal differentiation (15D). qRT-PCR expression was performed for other adipogenic genes *DNMT3A, DNMT3B, DNMT1, MBD1, MBD2, MBD3, MBD4, MeCP21, MeCP2, UHRF1, UHRF2, CBX5, ZFP467, TCF7L1, FOXC2, KLF2, VDR, FOXO1, DGAT1, CEBP□, KLF5, KLF6, STAT5a, EBF1, ZFP423, PDGFR□, PDGFRβl52, Pbx115, Fab4, Resistin, VEGFc, VEGFb, CD36, GLUT4, Scd1, SREBP1*that show inverse relation between methylation and gene expression as cells differentiate (Figure 6, FigureS4). The result showed threefold increase in *TET3* transcription increased during initial differentiation. However, *TET1* and *TET2* do not show any statistically significant change in expression[68]. We performed GO analyses using a list of genes with differentially methylated promoter regions[21][22][23]. Interestingly, the GO enrichment results aligned with our observations, suggesting that genes with hypomethylated promoters at 4H and 2D were associated with adipogenic processes-. At the same time, genes with hypermethylated promoters at 15D were enriched with GO terms related to white fat cell differentiation, brown fat cell differentiation, regulation of lipid metabolic processes, IL-7 signaling, TGF-β signaling pathway, DNA replication, insulin-like factors, regulation of localization, and adipogenesis genes (Figure 7A). Genes with hypomethylated promoters were identified in specific gene regions, indicating a link between DNA methylation reprogramming and preadipocyte differentiation. At 15D hyper-DMR, pathways associated with Lyz2, such as lysozyme activity and peptidoglycan muralytic activity, are upregulated. Upregulated expression of Lyz2 during 3T3L1 differentiation maintains the expression of genes associated with adipogensis and the differentiation of adipocytes[65]. This suggests that DNA methylation rapidly modifies gene loci following exposure to differentiation stimuli, eventually leading to gene expression variation.

**Figure 5:**
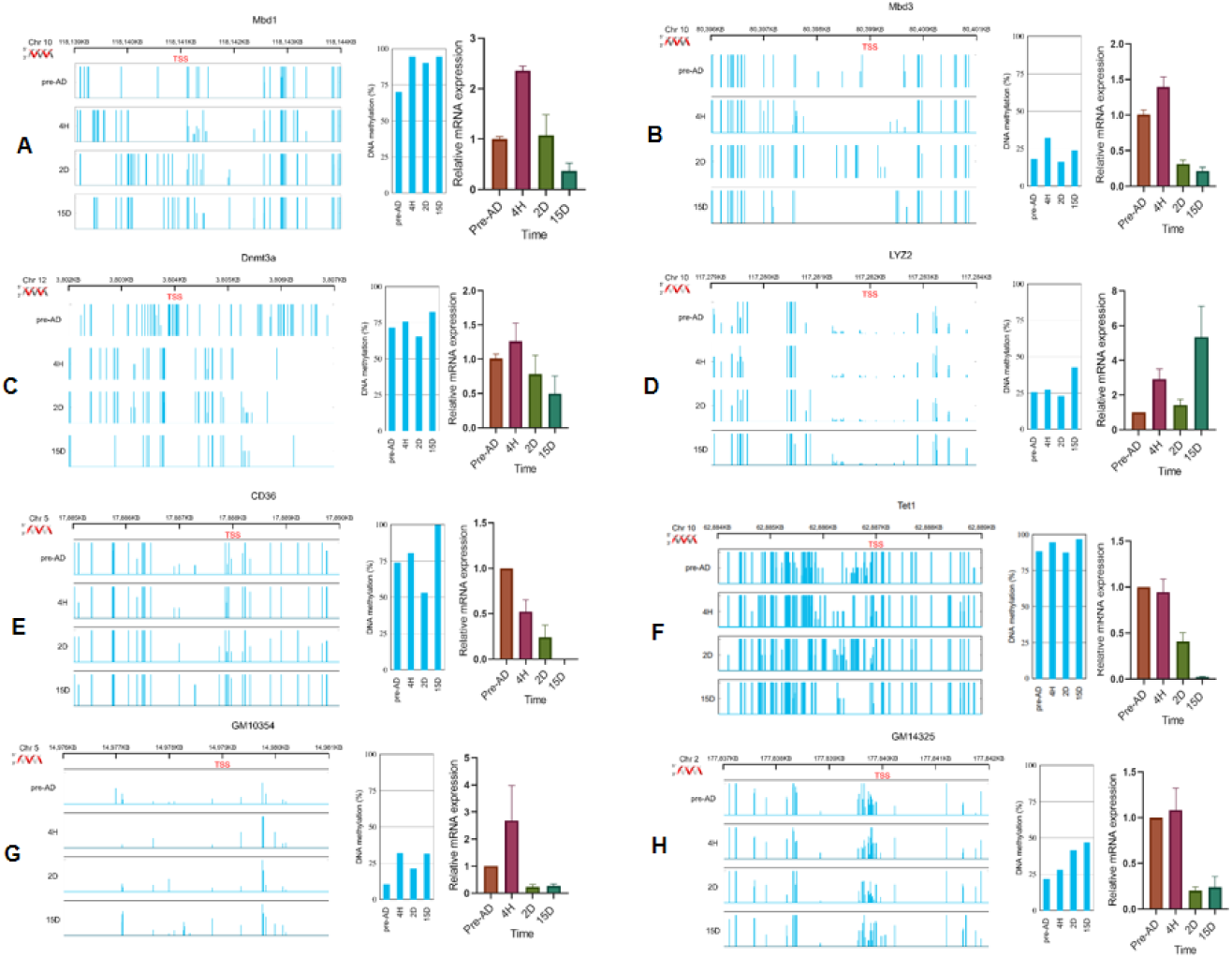
Illustrative presentation of a genome browser view displaying DNA methylation pattern in the promoters of genes A) MBD1, B) MBD3, C) DNMT3A, D) Lyz2, E) CD36, F) Tet1, G) GM10354, H) GM14325, and average DNA methylation level at the promoters of these genes along with genes expression profiles at indicated time points.

**Figure 6:**
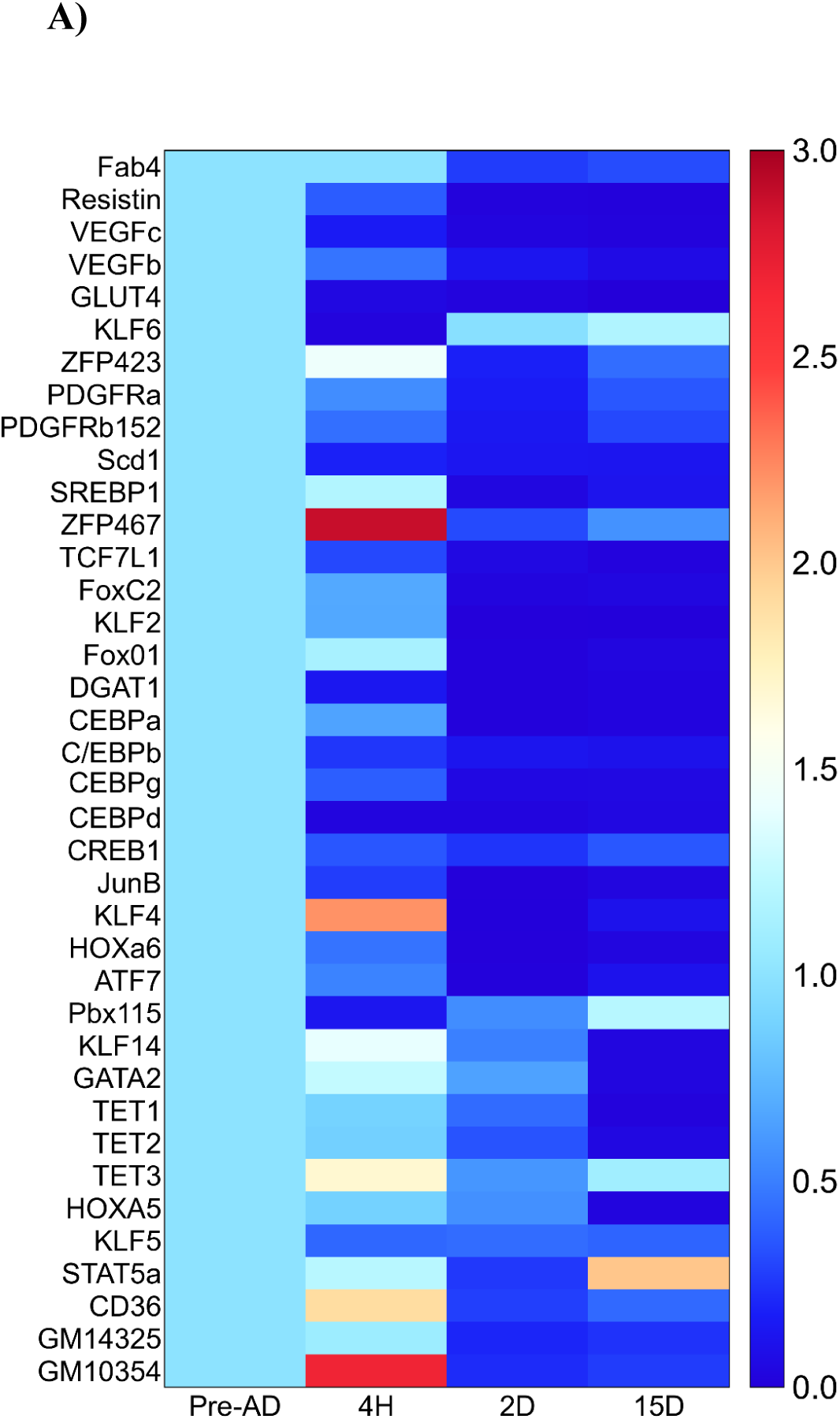
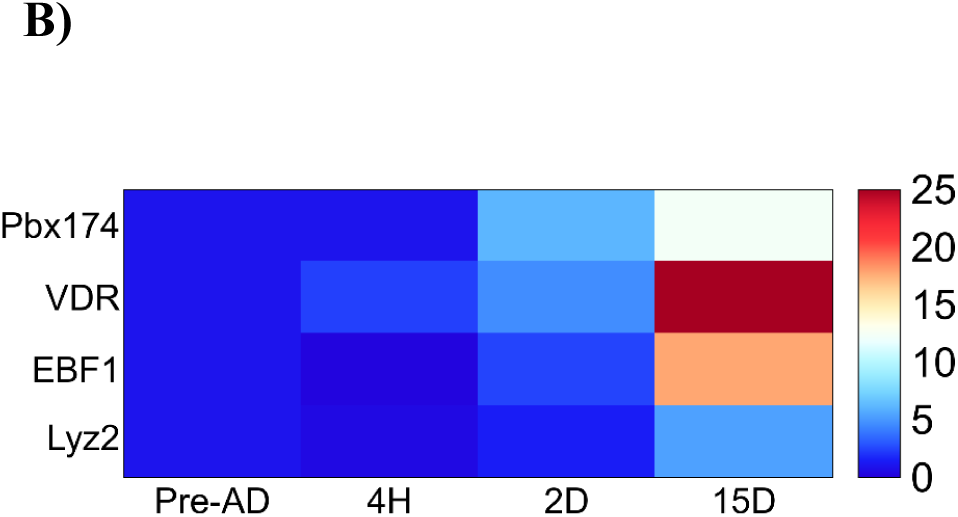
Using qRT-PCR, heatmaps of gene expression values depicting clustering of genes among undifferentiated and differentiated cells based on mRNAs expression for a set of genes (p-value<0.05). Samples are represented in rows, and genes are in columns. Low to high expression is given by blue to brown. A) Heatmap depicts the genes with up to 3-fold change in expression. B) Genes with more than a 5-fold change in expression.

**Figure 7:**
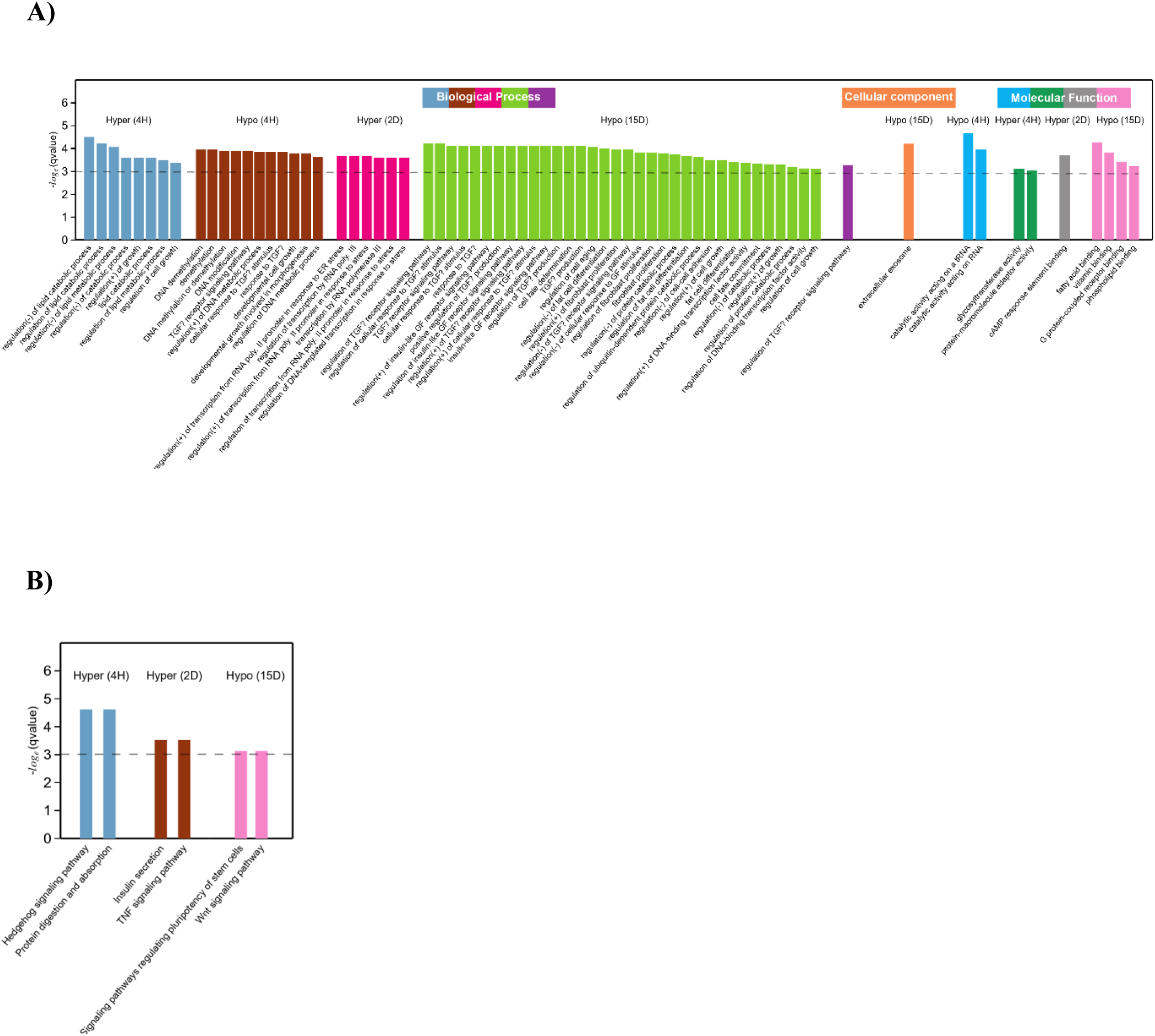
A) GO and B) KEGG pathway recruitment analysis of differentially expressed genes (DEGs) in the DMRs: undifferentiated and differentiated cells were categorized into various functional groups: Biological Process (BP), Cellular component (CC), and Molecular Function (MF).

### GO and KEGG analysis

The interplay between cell cycle regulators and differentiation factors triggers a chain of events leading to the adipocyte phenotype[26]. Adipogenesis is a multi-stage process characterized by a specific gene expression pattern[20], [27], [69], [70][71]. In the process of adipocyte differentiation from pluripotent stem cells, there are two distinct stages. The initial phase, termed determination, encompasses the commitment of pluripotent stem cells to preadipocytes. Preadipocytes may exhibit morphological similarities to their precursor cells but undergo a loss of versatility to differentiate into alternative cell types. In the subsequent phase, referred to as terminal differentiation, preadipocytes progressively assume the feature of mature adipocytes and attain functional capabilities, including lipid transport and synthesis, insulin sensitivity, and the secretion of adipocyte-specific proteins[20][27], [28][72]. During the terminal differentiation stage, there is a notable surge in the newly formed synthesis of fatty acids. Transcription factors collaborate with genes associated with adipocytes, working in tandem to maintain the progression of precursor adipocytes into fully mature adipocytes[31], [73], [74].

To comprehend the roles of DEGs (Differentially expressed genes), we conducted GO enrichment analysis to investigate their participation in biological processes, cellular components, and molecular functions (Figure 7A). DEGs were significantly enriched in metabolism-related pathways, including white cell differentiation, brown cell differentiation, transforming growth factor beta signalling, lipid metabolic pathways, interleukin signalling, DNA replication, fat cell differentiation, insulin-like growth factors, and steroid metabolic processes. We conducted a GO enrichment analysis to examine the biological significance of the DEGs in adipocytes. Among the biological processes, many DEGs were involved in cellular and metabolic processes. Most DEGs were associated with cell components and organelles in the cellular component category. Regarding molecular function, many DEGs were linked to catalytic activity and transcription regulator activity. This analysis helps to understand the functional roles and relationships of the DEGs in DMRs in adipocyte biology.

### KEGG pathway analysis of DEGs

We exploited the KEGG Pathway database to explore the signalling pathways associated with the DEGs in adipogenesis. This analysis revealed specific components that play a role in adipogenesis. Among the top 20 significant pathways of DEGs, the most prominent enrichments were observed in metabolic pathways, lipid biosynthesis, and the steroid metabolism pathway[43]. On the other hand, the TNF signalling pathway, hedgehog signalling pathway, insulin secretion, protein digestion and absorption, signalling pathways regulating pluripotency of stem cells, and Wnt signalling pathway were predominantly enriched (Figure 7B). This analysis provides valuable insights into the molecular mechanisms and biological processes involved in adipogenesis and helps to identify key pathways that may regulate the differentiation and function of adipocytes.

Identifying and verifying key genes regulating adipogenesis is crucial for understanding the molecular mechanisms underlying adipocyte differentiation. We compared DEGs in DMRs between the control and three experimental groups to identify potential candidate essential genes that regulate adipogenesis in 3T3-L1 cells[24], [25], [75]. This study reveals that groups’ differences in gene expression are linked to the differentiation of fat cells, lipid accumulation, and insulin production in treated 3T3-L1 cells. DNA methylation may also impact gene expression associated with these pathways.

### Dynamic DNA methylation at promoter of Lyz2

This study found that the Lyz2 gene’s promoter region was hypermethylated in terminally differentiated adipocytes (Figure S9). The CGs present at promoter region depicted substantial change in DNA methylation. Beyond that, CpG methylation was dynamic. Hypermethylation at promoter region at TD has been reported in some cases previously, and its causal role in disorders[76][77].

This is the first study to demarcate dynamic DNA hypermethylation at the promoter of Lyz2 gene during differentiation of preadipocytes indicating the causal role of DNA methylation at Lyz2 promoter that is deemed to be of great relevance.

### Non-CpG DNA methylation

The present study focused on variations in the methylome across the entire genome at the single CG dinucleotide resolution of preadipocytes and adipocytes. However, some non-CpG changes have been observed[78][79]. Variable non-CpG methylation, i.e., CHH and CHG, are observed. In the case of preadipocytes, the CHH and CHG were around 0.86 and 0.85, respectively, which subsequently decreased at 4H to 0.83 and 0.82, respectively. Cytosine methylation in non-CG sites further increased to 0.94 and 0.95 at 15 days, demonstrating the dynamic variations in non-CpG methylation as the differentiation of preadipocytes progressed (Figure S1). This aspect of preadipocyte differentiation may be further studied to establish the role of non-CpG methylation in pre-adipocyte differentiation.

## DISCUSSION

Emerging data indicates that cellular identity is also determined by a distinct DNA methylation pattern[44], [80], [81][62], [70], [82]. The function of DNA methylation in 3T3L1 cell differentiation has been investigated here, as the precise regulatory effects and underlying mechanisms on preadipocytes’ transformation into adipocytes are still unknown. To provide insight into these DNA methylation-mediated regulatory processes, we employed pre-adipocytes as a model system to see dynamic DNA methylation patterns as cells proceed to differentiate. Employing WGBS methodology, the study revealed the DNA methylation during different time periods. Previous studies have focused on earlier time periods ranging from a few hours to 6 days. To our knowledge, this is the first attempt to investigate high-resolution genome-wide DNA methylation at time intervals of 4H, 2D, and 15D, later correspond to the preadipocytes’ terminal differentiation.

Here, we have presented a comprehensive bisulfite sequencing workflow and analysis for 3T3L1 cells (Figure S2); at the early stages, i.e., during initial differentiation of pre-adipocytes, DNA methylation was found to be significantly reduced at CpG sites as well as non-CpG sites. However, as the differentiation progresses, the DNA methylation pattern is regained in the later stages, i.e., terminal differentiation. We also highlight varying global DNA methylation patterns and ‘hotspots’ site-specific variability. Transcription factor hotspots show cooperative binding of multiple transcription factors and restructure the chromatin structure within hours after induction of adipogenesis[29], [49], [82]. Analysis of CpG-rich hotspots on chr5 and chr8 and their DNA methylation status at four different time points suggests the importance of DNA methylation in gene regulatory functions. DMR analysis reduces the likelihood of negative random associations compared to single CpG site-based data[4], [83]. We found a subset of known and unknown DMRs in 3T3L1, which exhibited changes during the differentiation of preadipocytes. Novel DMRs with adipogenic genes were found. We discovered around 200, 492, and 1036 novel DMRs at 4H, 2D, and 15D. Furthermore, hypo-DMRs were pre-dominant at 4 H and 2D post-differentiation. At 15 days, the hyper-DMRs were enriched. Also, the idiogram of DMRs (Figure S5) confirmed that the distribution of DMRs was not uniform but was rather restricted to specific chromosomes. To further understand the role of DMRs in gene regulatory functions, hotspots overlapped with DMRs. In some cases, there was a partial overlap between hotspots and DMRs, whereas in others, complete overlap confirmed the correlation of gene regulatory functions with DMRs.

It is suggested that upregulation of DNA methylation at promoters leads to the repression of genes. On the contrary, DNA methylation at promoters can increase gene expression (this study and [72]). We found that DNA methylation at promoters of MBD1, MBD3, DNMT3A, LYZ2, CD36, TET1, GM10354, and GM14325 was upregulated at 15 days; however, the gene expression was decreased in all the cases except Lyz2,GM10354 and GM41325 where the DNA methylation and gene expression was positively correlated. At terminal differentiation, expression of VDR, EBF1, GM14325, Lyz2, GM10354 and C/EBPg was highly enhanced. Most of the adipogenic genes indicate decreased gene expression with increased promoter methylation, with few exceptions where an inverse relation is observed at initiation of terminal differentiation and terminal differentiation (Figure 8). Therefore, it is suggested that the relationship between promoter methylation and gene activity is complex and context-dependent. Further, key transcription factors and DEGs are significantly upregulated or downregulated, and the methylation status of key transcription factor promoters was altered. These findings provide valuable insights into regulating various adipocyte-specific genes during the process of adipogenesis.

**Figure 8:**
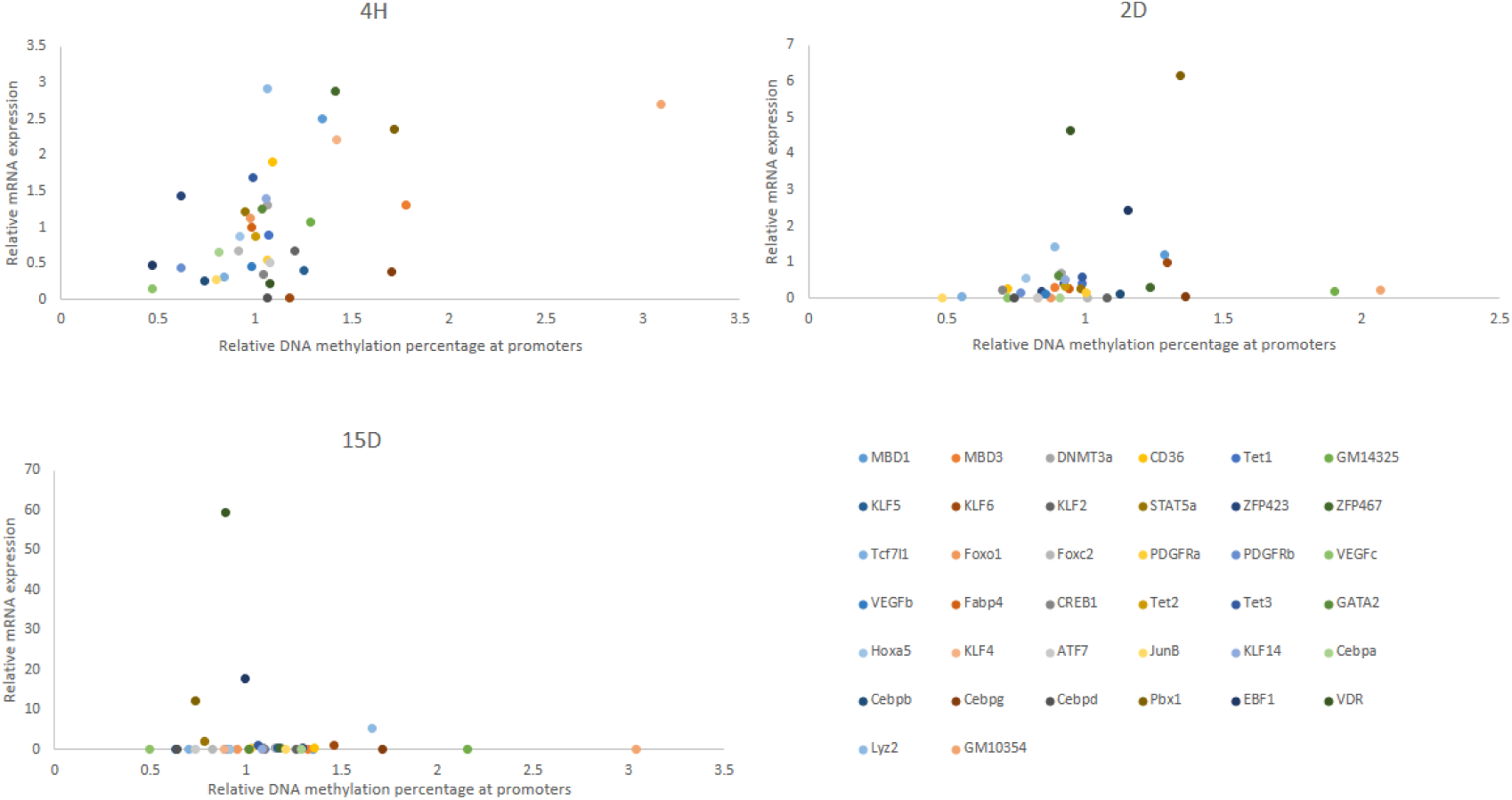
The connection between DNA methylation and gene expression was examined by plotting matched methylation and gene expression data obtained through Bisulfite sequencing of NIH 3T3L1 and qRT-PCR. The x-axis illustrates the extent of methylation at CG sites within the promoter, while the y-axis represents relative mRNA expression for the selected genes.

Furthermore, in order to ascertain the potential connection between alterations in DNA methylation and cellular functions, we conducted GO and KEGG analyses on genes exhibiting DMRs to examine enriched pathways. Genes featuring hypo-DMRs were found to be linked with the process of fat cell differentiation, DNA methylation and demethylation, cell fate determination, fibroblast proliferation, extracellular exosome, fatty acid binding, GPCR binding, and phospholipid binding. However, the hyper-DMRs were associated with lipid catabolic process and cAMP response element binding.

Our study has contributed to a deeper understanding of adipogenesis mechanisms while identifying potential epigenetic targets for regulating this process. The comparative analysis of hyper and hypo-DMRs at 2D and 15D (Table S5) signifies that at terminal differentiation some pathways are part of hyper-DMRs such as positive regulation of mitotic nuclear division, positive regulation of glucose metabolic process, glycogen biosynthetic process, glucan biosynthetic process whereas hypo-DMRs consist of pathways such as regulation of fat cell differentiation, regulation of protein localization to the nucleus, regulation of osteoblast differentiation.

Our study has demonstrated a dynamic DNA methylation pattern during pre-adipocyte differentiation. We have shown loss of DNA methylation at initial differentiation and regain of methylation pattern at the terminal stage of differentiation. This work has investigated the epigenetic mark DNA methylation associated with the differentiation of preadipocytes and its association with relevant genes associated with adipogenesis. The promoter of lyz2 showed increased methylation and gene expression at terminal differentiation.

In summary, cytosine methylation, as an epigenetic mechanism, regulates gene expression during adipogenesis. Understanding the dynamic changes in DNA methylation and its impact on adipocyte differentiation and function is crucial in unravelling the molecular mechanisms underlying obesity and related metabolic disorders.

## CONCLUSION

It is evident that DNA methylation plays a significant role in the lineage-specific development of adipocytes. Hypermethylation occurs during terminal differentiation, suggesting DNA methylation’s role in maintaining mature adipocytes. Few gene promoters, when hypermethylated, show high gene activity. Therefore, DNA methylation-dependent gene expression is context-dependent. Further understanding of the regulatory role of non-CpG methylation and targets for dedifferentiation can offer a more thorough insight into the epigenetic control involved in the differentiation of adipocytes.

## Supporting information

spdf

## Data and code availability

The raw Whole Genome Bisulphite Sequencing Data was deposited in the National Centre for Biotechnology Information Sequence Read Archive under Bioproject code **PRJNA1034485**. The *Mus musculus* mm10 data was used as the reference for alignment with raw data.

### Code availability

The method presented in the paper, and the code of this study will be shared via a readme file/Google Drive link. (https://drive.google.com/drive/folders/1bQ6v1cqduhcwVjTIsPfHN2WyskgRXVmu?usp=drive_link)

## Author Contributions

BY and VR conceived and designed the study. BY performed the wet lab experiments. BY and DS analysed the data and contributed equally. DS helped with bioinformatic analysis. VR and SM supervised and reviewed the study. BY, DS and VR wrote the manuscript. VR and SM edited the manuscript.

## Acknowledgements

We thank the Executive director, National Agri-Food Biotechnology Institute (NABI), Mohali for research facilities and Department of Biotechnology (DBT), New Delhi for research funding. BY acknowledges the financial assistance from DBT in the form of JRF and SRF. The authors thank the National Agri-Food Biotechnology Institute of the Department of Biotechnology (DBT), Ministry of Science and Technology, Government of India, for supporting high-performance computing. We acknowledge National supercomputing Mission (NSM) for providing computing resources of ‘PARAM Smriti’ at NABI, Mohali, which is implemented by C-DAC and supported by Ministry of Electronics and Information Technology (MeitY) and Department of Science and Technology (DST), Government of India.

## References

[1] R. Holliday, “Epigenetics: a historical overview,” Epigenetics, vol. 1, no. 2, pp. 76–80, 2006.

[2] M. M. Suzuki and A. Bird, “DNA methylation landscapes: provocative insights from epigenomics,” Nat. Rev. Genet., vol. 9, no. 6, pp. 465–476, 2008.

[3] S. Yeo, S. Jeong, J. Kim, J.-S. Han, Y.-M. Han, and Y.-K. Kang, “Characterization of DNA methylation change in stem cell marker genes during differentiation of human embryonic stem cells,” Biochem. Biophys. Res. Commun., vol. 359, no. 3, pp. 536– 542, 2007.

[4] A. Roy, S. S. Padhi, I. Khyriem, S. Nikose, and R. S. Bharathavikru, “Resetting the epigenome: Methylation dynamics in cancer stem cells,” Front. Cell Dev. Biol., p. 1934, 2022.

[5] R. Singal and G. D. Ginder, “DNA methylation,” Blood, J. Am. Soc. Hematol., vol. 93, no. 12, pp. 4059–4070, 1999.

[6] L. D. Moore, T. Le, and G. Fan, “DNA methylation and its basic function,” Neuropsychopharmacology, vol. 38, no. 1, pp. 23–38, 2013.

[7] P. M. Das and R. Singal, “DNA methylation and cancer,” J. Clin. Oncol., vol. 22, no. 22, pp. 4632–4642, 2004.

[8] R. Das et al., “Computational prediction of methylation status in human genomic sequences,” Proc. Natl. Acad. Sci., vol. 103, no. 28, pp. 10713–10716, 2006.

[9] M. Kulis and M. Esteller, “DNA methylation and cancer,” Adv. Genet., vol. 70, pp. 27–56, 2010.

[10] Y. Saito et al., “Increased protein expression of DNA methyltransferase (DNMT) 1 is significantly correlated with the malignant potential and poor prognosis of human hepatocellular carcinomas,” Int. J. cancer, vol. 105, no. 4, pp. 527–532, 2003.

[11] K. D. Robertson and P. A. Jones, “DNA methylation: past, present and future directions,” Carcinogenesis, vol. 21, no. 3, pp. 461–467, 2000.

[12] K. D. Robertson, “DNA methylation and human disease,” Nat. Rev. Genet., vol. 6, no. 8, pp. 597–610, 2005.

[13] Z. D. Smith and A. Meissner, “DNA methylation: roles in mammalian development,” Nat. Rev. Genet., vol. 14, no. 3, pp. 204–220, 2013.

[14] H. Cedar, “DNA methylation and gene activity.,” Cell, vol. 53, no. 1, pp. 3–4, 1988.

[15] N. Petryk, S. Bultmann, T. Bartke, and P.-A. Defossez, “Staying true to yourself: mechanisms of DNA methylation maintenance in mammals,” Nucleic Acids Res., vol. 49, no. 6, pp. 3020–3032, 2021.

[16] M. Weber et al., “Distribution, silencing potential and evolutionary impact of promoter DNA methylation in the human genome,” Nat. Genet., vol. 39, no. 4, pp. 457–466, 2007.

[17] P. A. Jones and D. Takai, “The role of DNA methylation in mammalian epigenetics,” Science (80-. )., vol. 293, no. 5532, pp. 1068–1070, 2001.

[18] P. A. Jones, “The DNA methylation paradox,” Trends Genet., vol. 15, no. 1, pp. 34– 37, 1999.

[19] P. A. Jones, “Functions of DNA methylation: islands, start sites, gene bodies and beyond,” Nat. Rev. Genet., vol. 13, no. 7, pp. 484–492, 2012.

[20] J.-E. Lee, H. Schmidt, B. Lai, and K. Ge, “Transcriptional and Epigenomic Regulation of Adipogenesis,” Mol. Cell. Biol., vol. 39, no. 11, pp. 1–20, 2019.

[21] K. E. Pinnick and F. Karpe, “DNA methylation of genes in adipose tissue,” Proc. Nutr. Soc., vol. 70, no. 1, pp. 57–63, 2011.

[22] H. Xie et al., “DNA methylation modulates aging process in adipocytes,” Aging Dis., vol. 13, no. 2, p. 433, 2022.

[23] G. Agha, E. A. Houseman, K. T. Kelsey, C. B. Eaton, S. L. Buka, and E. B. Loucks, “Adiposity is associated with DNA methylation profile in adipose tissue,” Int. J. Epidemiol., vol. 44, no. 4, pp. 1277–1287, 2015.

[24] K. Sarjeant and J. M. Stephens, “Adipogenesis,” Cold Spring Harb. Perspect. Biol., vol. 4, no. 9, pp. a008417–a008417, 2012.

[25] M. I. Lefterova and M. A. Lazar, “New developments in adipogenesis,” Trends Endocrinol. Metab., vol. 20, no. 3, pp. 107–114, 2009.

[26] E. D. Rosen and B. M. Spiegelman, “Molecular regulation of adipogenesis,” Annu. Rev. Cell Dev. Biol., vol. 16, no. 1, pp. 145–171, 2000.

[27] E. D. Rosen, C. J. Walkey, P. Puigserver, and B. M. Spiegelman, “Transcriptional regulation of adipogenesis,” Genes Dev., vol. 14, no. 11, pp. 1293–1307, 2000.

[28] R. Siersbæk, R. Nielsen, and S. Mandrup, “Transcriptional networks and chromatin remodeling controlling adipogenesis,” Trends Endocrinol. Metab., vol. 23, no. 2, pp. 56–64, 2012.

[29] R. Siersbæk et al., “Extensive chromatin remodelling and establishment of transcription factor ‘hotspots’ during early adipogenesis,” EMBO J., vol. 30, no. 8, pp. 1459–1472, 2011.

[30] T. S. Mikkelsen et al., “Comparative epigenomic analysis of murine and human adipogenesis,” Cell, vol. 143, no. 1, pp. 156–169, 2010.

[31] R. Siersbæk et al., “Dynamic rewiring of promoter-anchored chromatin loops during adipocyte differentiation,” Mol. Cell, vol. 66, no. 3, pp. 420–435, 2017.

[32] M. Aso et al., “First-in-human autologous implantation of genetically modified adipocytes expressing LCAT for the treatment of familial LCAT deficiency,” Heliyon, vol. 8, no. 11, 2022.

[33] A. M. Bolger, M. Lohse, and B. Usadel, “Trimmomatic: a flexible trimmer for Illumina sequence data,” Bioinformatics, vol. 30, no. 15, pp. 2114–2120, 2014.

[34] F. Krueger and S. R. Andrews, “Bismark: a flexible aligner and methylation caller for Bisulfite-Seq applications,” bioinformatics, vol. 27, no. 11, pp. 1571–1572, 2011.

[35] A. Akalin et al., “methylKit: a comprehensive R package for the analysis of genome-wide DNA methylation profiles,” Genome Biol., vol. 13, no. 10, pp. 1–9, 2012.

[36] H. Wu et al., “Detection of differentially methylated regions from whole-genome bisulfite sequencing data without replicates,” Nucleic Acids Res., vol. 43, no. 21, pp. e141--e141, 2015.

[37] G. Yu, L.-G. Wang, and Q.-Y. He, “ChIPseeker: an R/Bioconductor package for ChIP peak annotation, comparison and visualization,” Bioinformatics, vol. 31, no. 14, pp. 2382–2383, 2015.

[38] K. Williams et al., “TET1 and hydroxymethylcytosine in transcription and DNA methylation fidelity,” Nature, vol. 473, no. 7347, pp. 343–348, 2011.

[39] R. Jaura et al., “Extended intergenic DNA contributes to neuron-specific expression of neighboring genes in the mammalian nervous system,” Nat. Commun., vol. 13, no. 1, pp. 1–18, 2022.

[40] Y. Xi et al., “Multi-omic characterization of genome-wide abnormal DNA methylation reveals diagnostic and prognostic markers for esophageal squamous-cell carcinoma,” Signal Transduct. Target. Ther., vol. 7, no. 1, pp. 1–13, 2022.

[41] F. Ram\’\irez et al., “deepTools2: a next generation web server for deep-sequencing data analysis,” Nucleic Acids Res., vol. 44, no. W1, pp. W160--W165, 2016.

[42] A. J. Bewick et al., “On the origin and evolutionary consequences of gene body DNA methylation,” Proc. Natl. Acad. Sci., vol. 113, no. 32, pp. 9111–9116, 2016.

[43] J. Park et al., “Targeted erasure of DNA methylation by TET3 drives adipogenic reprogramming and differentiation,” Nat. Metab., vol. 4, no. 7, pp. 918–931, 2022.

[44] H. Sakamoto et al., “Sequential changes in genome-wide DNA methylation status during adipocyte differentiation,” Biochem. Biophys. Res. Commun., vol. 366, no. 2, pp. 360–366, 2008.

[45] C. Vinson and R. Chatterjee, “CG methylation,” Epigenomics, vol. 4, no. 6, pp. 655– 663, 2012.

[46] A. Doi et al., “Differential methylation of tissue-and cancer-specific CpG island shores distinguishes human induced pluripotent stem cells, embryonic stem cells and fibroblasts,” Nat. Genet., vol. 41, no. 12, pp. 1350–1353, 2009.

[47] R. Fenouil et al., “CpG islands and GC content dictate nucleosome depletion in a transcription-independent manner at mammalian promoters,” Genome Res., vol. 22, no. 12, pp. 2399–2408, 2012.

[48] R. Straussman et al., “Developmental programming of CpG island methylation profiles in the human genome,” Nat. Struct. Mol. Biol., vol. 16, no. 5, pp. 564–571, 2009.

[49] R. Siersbæk et al., “Transcription factor cooperativity in early adipogenic hotspots and super-enhancers,” Cell Rep., vol. 7, no. 5, pp. 1443–1455, 2014.

[50] J. D. Cleary and C. E. Pearson, “The contribution of cis-elements to disease-associated repeat instability: clinical and experimental evidence,” Cytogenet. Genome Res., vol. 100, no. 1–4, pp. 25–55, 2003.

[51] P.-M. Challita, D. Skelton, A. El-Khoueiry, X.-J. Yu, K. Weinberg, and D. B. Kohn, “Multiple modifications in cis elements of the long terminal repeat of retroviral vectors lead to increased expression and decreased DNA methylation in embryonic carcinoma cells,” J. Virol., vol. 69, no. 2, pp. 748–755, 1995.

[52] T. J. Peters et al., “De novo identification of differentially methylated regions in the human genome,” Epigenetics Chromatin, vol. 8, pp. 1–16, 2015.

[53] F. Song et al., “Association of tissue-specific differentially methylated regions (TDMs) with differential gene expression,” Proc. Natl. Acad. Sci., vol. 102, no. 9, pp. 3336–3341, 2005.

[54] D. Beck, M. Ben Maamar, and M. K. Skinner, “Genome-wide CpG density and DNA methylation analysis method (MeDIP, RRBS, and WGBS) comparisons,” Epigenetics, vol. 17, no. 5, pp. 518–530, 2022.

[55] R. G. Smith et al., “A meta-analysis of epigenome-wide association studies in Alzheimer’s disease highlights novel differentially methylated loci across cortex,” Nat. Commun., vol. 12, no. 1, p. 3517, 2021.

[56] L. Zhang et al., “Epigenome-wide meta-analysis of DNA methylation differences in prefrontal cortex implicates the immune processes in Alzheimer’s disease,” Nat. Commun., vol. 11, no. 1, p. 6114, 2020.

[57] P. A. Jones and S. B. Baylin, “The fundamental role of epigenetic events in cancer,” Nat. Rev. Genet., vol. 3, no. 6, pp. 415–428, 2002.

[58] D. Aran, G. Toperoff, M. Rosenberg, and A. Hellman, “Replication timing-related and gene body-specific methylation of active human genes,” Hum. Mol. Genet., vol. 20, no. 4, pp. 670–680, 2011.

[59] S. A. Bert et al., “Regional activation of the cancer genome by long-range epigenetic remodeling,” Cancer Cell, vol. 23, no. 1, pp. 9–22, 2013.

[60] C. G. Spilianakis, M. D. Lalioti, T. Town, G. R. Lee, and R. A. Flavell, “Interchromosomal associations between alternatively expressed loci,” Nature, vol. 435, no. 7042, pp. 637–645, 2005.

[61] L. E. Docherty et al., “Genome-wide DNA methylation analysis of patients with imprinting disorders identifies differentially methylated regions associated with novel candidate imprinted genes,” J. Med. Genet., vol. 51, no. 4, pp. 229–238, 2014.

[62] W. He et al., “Defining differentially methylated regions specific for the acquisition of pluripotency and maintenance in human pluripotent stem cells via microarray,” PLoS One, vol. 9, no. 9, p. e108350, 2014.

[63] Q. Ma et al., “Specific hypomethylation programs underpin B cell activation in early multiple sclerosis,” Proc. Natl. Acad. Sci., vol. 118, no. 51, p. e2111920118, 2021.

[64] J. Park et al., “Functional methylome analysis of human diabetic kidney disease,” JCI insight, vol. 4, no. 11, 2019.

[65] A. Lluch, J. Latorre, J. M. Fernández-Real, and J. M. Moreno-Navarrete, “Lysozyme Gene Expression in 3T3-L1 Cells Sustains Expression of Adipogenic Genes and Adipocyte Differentiation,” Front. Cell Dev. Biol., vol. 10, p. 914788, 2022.

[66] K. Chechi, Y. Gelinas, P. Mathieu, Y. Deshaies, and D. Richard, “Validation of reference genes for the relative quantification of gene expression in human epicardial adipose tissue,” PLoS One, vol. 7, no. 4, p. e32265, 2012.

[67] T. D. Schmittgen and K. J. Livak, “Analyzing real-time PCR data by the comparative CT method,” Nat. Protoc., vol. 3, no. 6, pp. 1101–1108, 2008.

[68] B. C. Jung et al., “TET3 plays a critical role in white adipose development and diet-induced remodeling,” Cell Rep., vol. 42, no. 10, 2023.

[69] M. Kulis, A. C. Queirós, R. Beekman, and J. I. Martín-Subero, “Intragenic DNA methylation in transcriptional regulation, normal differentiation and cancer,” Biochim. Biophys. Acta (BBA)-Gene Regul. Mech., vol. 1829, no. 11, pp. 1161–1174, 2013.

[70] P. Madrigal et al., “Epigenetic and transcriptional regulations prime cell fate before division during human pluripotent stem cell differentiation,” Nat. Commun., vol. 14, no. 1, p. 405, 2023.

[71] J. Park et al., “Targeted erasure of DNA methylation by TET3 drives adipogenic reprogramming and differentiation,” Nat. Metab., vol. 4, no. 7, pp. 918–931, 2022.

[72] V. Rishi et al., “CpG methylation of half-CRE sequences creates C/EBPα binding sites that activate some tissue-specific genes,” Proc. Natl. Acad. Sci., vol. 107, no. 47, pp. 20311–20316, 2010.

[73] Y. C. Lim, S. Y. Chia, S. Jin, W. Han, C. Ding, and L. Sun, “Dynamic DNA methylation landscape defines brown and white cell specificity during adipogenesis,” Mol. Metab., vol. 5, no. 10, pp. 1033–1041, 2016.

[74] L. Laurent et al., “Dynamic changes in the human methylome during differentiation,” Genome Res., vol. 20, no. 3, pp. 320–331, 2010.

[75] Q. Q. Tang and M. D. Lane, “Adipogenesis: from stem cell to adipocyte,” Annu. Rev. Biochem., vol. 81, pp. 715–736, 2012.

[76] R. A. Zorzo et al., “LDLR gene’s promoter region hypermethylation in patients with familial hypercholesterolemia,” Sci. Rep., vol. 13, no. 1, p. 9241, 2023.

[77] J. Latorre et al., “Adipose tissue knockdown of lysozyme reduces local inflammation and improves adipogenesis in high-fat diet-fed mice,” Pharmacol. Res., vol. 166, p. 105486, 2021.

[78] A. S. Dey, “Functional Analysis of TET-Family 5-Methylcytosine Dioxygenases in Non-CpG DNA Demethylation.” University of Missouri-Kansas City, 2023.

[79] D. Ramasamy, A. K. D. M. Rao, T. Rajkumar, and S. Mani, “Experimental and Computational Approaches for Non-CpG Methylation Analysis,” Epigenomes, vol. 6, no. 3, p. 24, 2022.

[80] M. Suelves, E. Carrió, Y. Núñez-Álvarez, and M. A. Peinado, “DNA methylation dynamics in cellular commitment and differentiation,” Brief. Funct. Genomics, vol. 15, no. 6, pp. 443–453, 2016.

[81] M. Berdasco and M. Esteller, “DNA methylation in stem cell renewal and multipotency,” Stem Cell Res. Ther., vol. 2, pp. 1–9, 2011.

[82] R. Lister et al., “Hotspots of aberrant epigenomic reprogramming in human induced pluripotent stem cells,” Nature, vol. 471, no. 7336, pp. 68–73, 2011.

[83] M. P. Campagna et al., “Epigenome-wide association studies: current knowledge, strategies and recommendations,” Clin. Epigenetics, vol. 13, no. 1, pp. 1–24, 2021.

