## Supplementary material for "WGBS of Differentiating Adipocytes Reveals Variations in DMRs and Context-Dependent Gene Expression": spdf

List of supplemental items:

**Figure S1**: Summary of WGBS sequencing read counts of undifferentiated pre-adipocytes (Pre-AD) and differentiated (4H, 2D, and 15D post differentiation).

**Figure S2**: Schematic diagram of the steps for Whole genome Bisulphite Sequencing (WGBS) and pipeline for data analysis, and associated data analysis programs.

**Figure S3**: Chromosome-wise DNA methylation pattern depicting various time points during pre-adipocyte differentiation.

**Figure S4**: qRT-PCR analysis: Bar graphs showing qRT-PCR analysis of various adipogenic genes such as DNMT3A, DNMT3B, DNMT1, MBD1, MBD2, MBD3, MBD4, MeCP21, MeCP2 2, UHRF1, UHRF2, CBX5.

**Figure S5**: Idiograms depicting the DMRs distribution on chromosomes. A) Hyper-DMRs at 15D, B) Hyper-DMRs at 2D, C) Hyper-DMRs at 4H, D) Hypo-DMRs at 15D, E) Hypo-DMRs at 2D, F) Hypo-DMRs at 4H.

**Figure S6**: Pearson Correlation between NABI control vs. AR control. Total locations of NABI control with coverage 10:29584. Total locations of AR pre-adipocyte with coverage 10: 32860286. Common locations used for correlation analysis: 28973.

**Figure S7**: Comparison of NABI & AR data. Red color indicates pre-adipocytes and green represents the adipocytes whereas blue indicates NABI control.

**Figure S8**: Correlation analysis at various depths (from 4 to 12) for NABI control vs AR control.

**Figure S9:** Hypermethylation at Lyz2 promoter at terminally differentiated adipocytes.

**Table S1**: List of Oligos used in the study for performing qRT-PCR to check adipogenic gene expression.

**Table S2: GO pathway analysis.** GO enrichment analysis of genes whose promoter overlapped with DMRs. For GO analysis, genes were grouped into different functional categories: biological process (BP), Cellular component (CC), and Molecular Function (MF). **KEGG pathway analysis** enrichKEGG of clusterProfiler is used for pathway analysis with BH procedure and (final parametric values will be after results are fixed pvalueCutoff = 1,pAdjustMethod = "BH",minGSSize = 1, maxGSSize = 500,qvalueCutoff = 1)

(https://docs.google.com/spreadsheets/d/13U_tKgUZ1KAUxRFsnBY8WN7UT46Z6ixS/edit?usp=sharing&ouid=106034465131344413756&rtpof=true&sd=true )

**Table S3**: Summary of AR WGBS dataset and read counts in pre-adipocytes and adipocytes depicting increased methylated CpGs as the pre-adipocytes differentiated. A) Mapping of raw reads with reference genome mm10. B) Total analysis of methylated cytosine showing CG cytosines and non-CG cytosines.

**Table S4**: Correlation analysis at various depths (from 4 to 12) for NABI control vs AR control.

**Table S5:** Comparison of methylation-dependent gene expression analysis of DMRs present at 2D and 15D; 15D depicting increased expression of certain pathways involved in dedifferentiation.

**Comparative analysis between the AR (Adipogenic reprogramming) dataset vs. the NABI dataset**

The NABI Whole genome bisulphite sequencing data referred to as the NABI dataset in the subsequent text was compared with the Adipogenic reprogramming dataset (AR dataset,[1]) to validate the results for the analysis of data in terms of whole genome and for comparison of NABI Sanger sequencing data of NIH 3T3L1. The total methylated CpG percentage varies by around 2-4% in completely differentiated adipocytes compared to pre-adipocytes (**Figure S1 & Table S3**) confirmed by both datasets. The bisulphite conversion rate was approximately the same in both cases. The mapping rate was also the same i.e. around 74% in all the cases except 2 days, around 85% (**FigureS1C & TableS3A**).

**Correlation analysis between AR and NABI dataset**

The evidence from the AR dataset suggests that unique DNA methylation establishes cellular identity, and our study supports this notion. In both studies, WGBS is used to investigate the dynamics of DNA methylation during adipocyte differentiation. The correlation between AR dataset and NABI dataset control is also evident which is 0.94 (**Figure S6**). At the early stage of adipogenesis, DNA methylation was specifically decreased at adipogenesis-related genomic loci which is quite evident in both the studies. The circos plot (**Figure S7**) depicts the chromosome-wise overlap.

The high correlation between AR and NABI data validates the significance of NABI data. The higher the depth, the higher the correlation was found (Supplementary Figure 3).

Given the resemblances we found between both the datasets, suggest DNA methylation plays a major regulatory mechanism during the differentiation of preadipocytes.

**A**

| Sample | Raw Reads | Clean Reads | Base quality Q20 (%) | Base quality Q30 (%) | GC (%) | Reads having 50% bases with quality <20 (%) | Reads having 50% bases with quality < 30 (%) |
|---|---|---|---|---|---|---|---|
| Pre-AD | 55,438,326 | 53,411,178 | 96.55 | 89.94 | 20.08 | 0 | 0.15 |
| 4H | 51,941,175 | 42,412,105 | 97.33 | 91.54 | 20.16 | 0 | 0.32 |
| 2D | 55,463,283 | 53,311,291 | 96.38 | 89.59 | 21.14 | 0 | 0.14 |
| 15D | 52,949,471 | 49,723,797 | 95.39 | 87.47 | 20.04 | 0 | 0.17 |

**B**

| | Pre-AD | 4H | 2D | 15D |
|---|---|---|---|---|
| Number of Reads aligned to Lambda Genome after de-duplication removal | 522,924 | 420,562 | 432,252 | 657,646 |
| Total Cytosines analysed | 4,288,500 | 3,488,969 | 3,612,914 | 5,080,239 |
| Total methylated Cytosines | 30,682 | 23,772 | 24,187 | 38,216 |
| Total unmethylated Cytosines | 4,257,818 | 3,465,197 | 3,588,727 | 5,042,023 |
| Bisulfite Conversion Efficiency (%) | 99.285 | 99.319 | 99.331 | 99.248 |

**C**

| Sample | Total Reads | Mapped Reads | Mapping Rate (%) | Duplicated reads | De-duplication rate (%) | Reads for methylation extraction |
|---|---|---|---|---|---|---|
| Pre-AD | 53,411,178 | 39,346,076 | 73.66 | 7,821,993 | 19.88 | 31,523,935 |
| 4H | 42,412,105 | 31,456,049 | 74.17 | 5,687,201 | 18.08 | 25,768,684 |
| 2Days | 53,311,291 | 45,153,417 | 84.70 | 8,364,937 | 18.53 | 36,788,357 |
| 15Days | 49,723,797 | 36,544,292 | 73.49 | 8,580,592 | 23.48 | 27,963,323 |

**D**

| Sample | Total no. of Cs analysed | Methylated Cs in CpG context | Unmethylated Cs in CpG context | mCpG (%) | mCHG (%) | mCHH (%) |
|---|---|---|---|---|---|---|
| Pre-AD | 1,608,609,236 | 41,240,654 | 27,786,970 | 59.75 | 0.86 | 0.85 |
| 4H | 1,319,747,325 | 33,756,587 | 22,922,625 | 59.60 | 0.83 | 0.82 |
| 2Days | 1,862,473,519 | 47,729,004 | 31,668,170 | 60.11 | 0.82 | 0.81 |
| 15Days | 1,425,151,215 | 38,393,366 | 22,286,277 | 63.27 | 0.94 | 0.95 |

**Figure S1**: Summary of WGBS sequencing read counts of undifferentiated pre-adipocytes(Pre-AD) and differentiated (4H, 2D, and 15D post differentiation). Bismark was used for paired-end mapping. Duplicated reads were removed using the deduplicate_bismark command. The genome-wide cytosine analysis was performed using the remaining reads. The methylation bias in the reads was determined with the –mbias option of Bismark Methylation Extractor; consequently, methylated CpGs were extracted**.** **(A)** Raw reads and their cleaning with Trimmomatic **(B)** Clean data was mapped to mouse genome mm10 by Bismark pipeline. Average coverage > 1x at each cytosine site. **(C)** Mapping of raw reads with reference genome mm10. **(D)** Total analysis of methylated cytosine showing CG cytosines and non-CG cytosines.

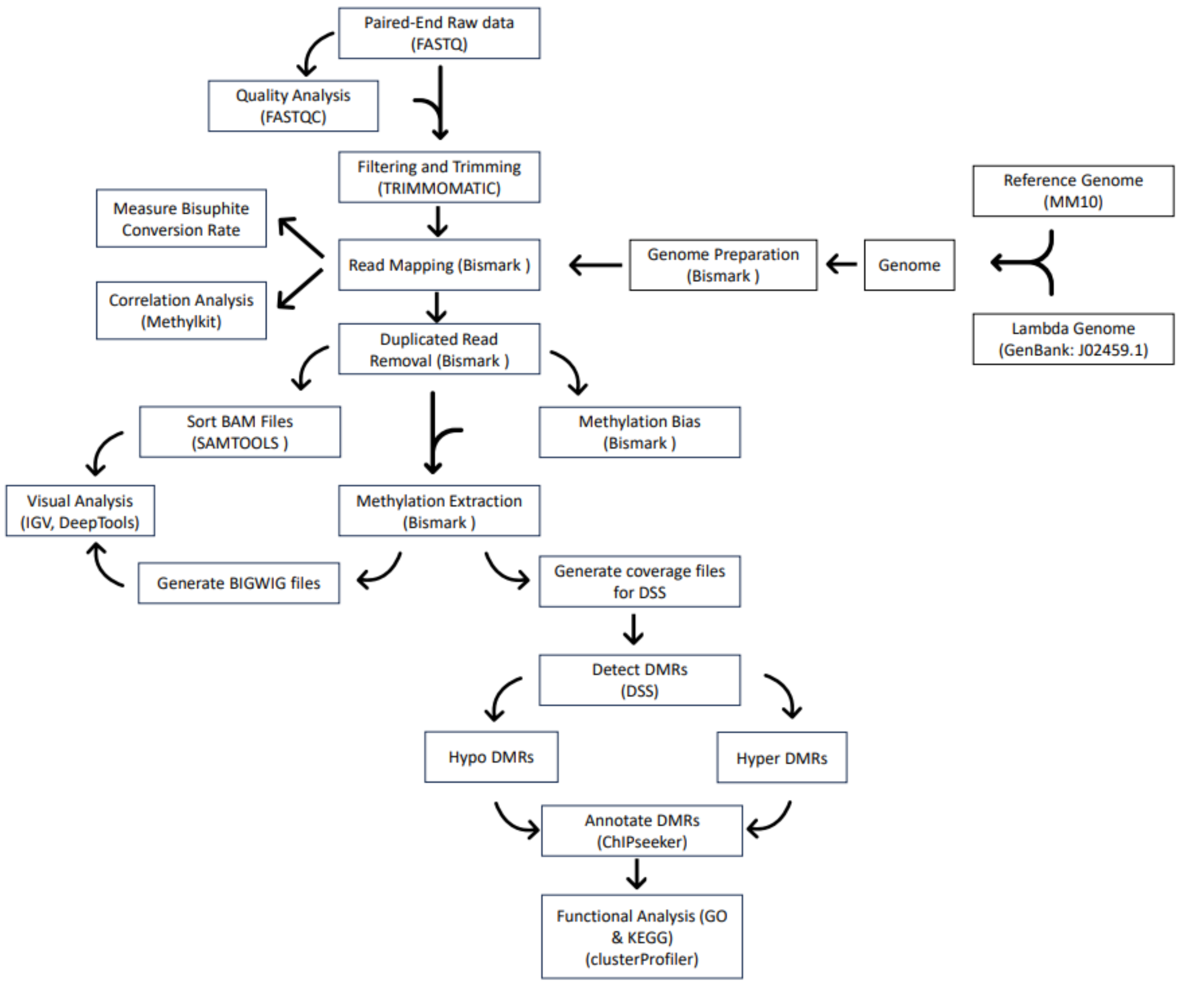


**Figure S2**: Schematic diagram of the steps for Whole genome Bisulphite Sequencing (WGBS) and pipeline for data analysis, and associated data analysis programs. FASTQ files generated from raw WGBS data were aligned with reference genome using Bismark. Output bam files were further processed through the remainder of the workflow pipeline.

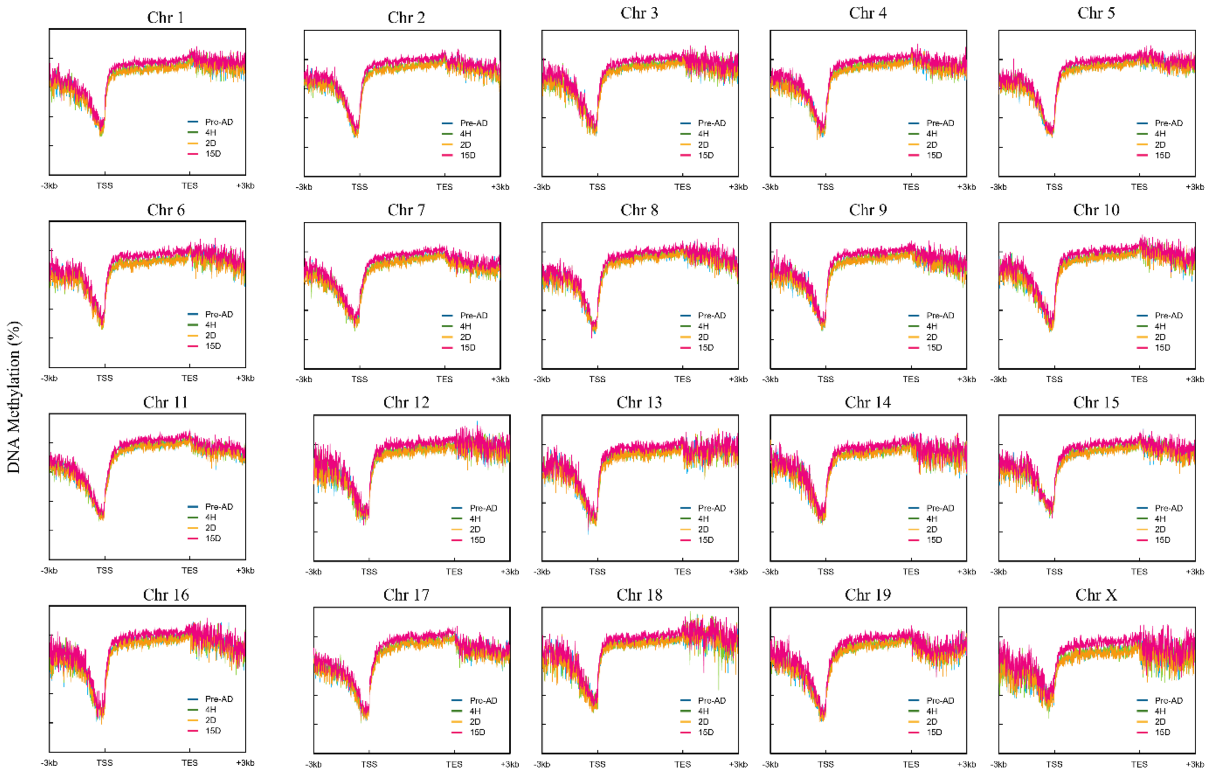


**Figure S3**: Chromosome-wise DNA methylation pattern depicting various time points during pre-adipocyte differentiation to mature adipocytes. X-axis: DNA methylation percentage. Y-axis- depicts Transcription start site (TSS), Transcription end site (TES), gene body, and promoter region.

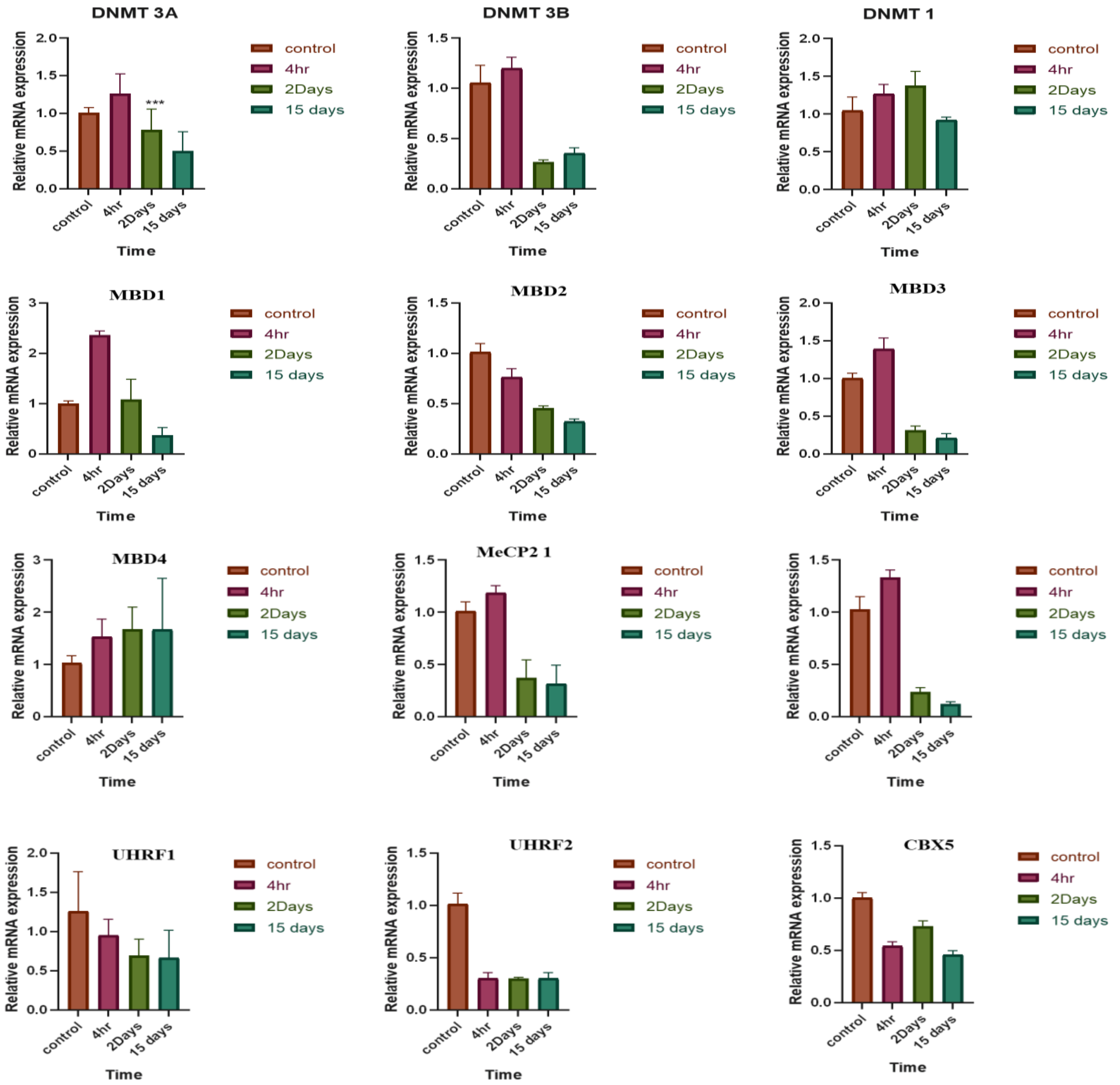


**Figure S4:** qRT-PCR analysis: Bar graphs showing qRT-PCR analysis of various adipogenic genes such as DNMT3A, DNMT3B, DNMT1, MBD1, MBD2, MBD3, MBD4, MeCP21, MeCP2 2, UHRF1, UHRF2, CBX5.

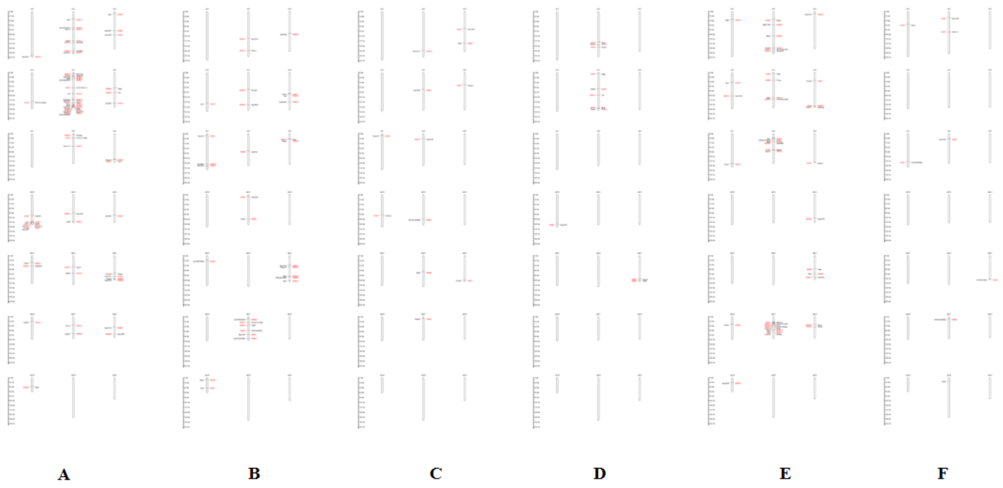


**Figure S5**: Idiograms depicting the DMRs distribution on chromosomes. A) Hyper-DMRs at 15D, B) Hyper-DMRs at 2D, C) Hyper-DMRs at 4H, D) Hypo-DMRs at 15D, E) Hypo-DMRs at 2D, F) Hypo-DMRs at 4H.

**Table S1:** Primers used for qRT-PCR.

| Adipogenic Genes | | |
|---|---|---|
| KLF5 | Forward Primer | GGCTCTCCCCGAGTTCACTA |
| | Reverse Primer | ATTACTGCCGTCTGGTTTGTC |
| KLF6 | Forward Primer | GTTTCTGCTCGGACTCCTGAT |
| | Reverse Primer | TTCCTGGAAGATGCTACACATTG |
| STAT5a | Forward Primer | CAGCCGTGGGATGCTATTGA |
| | Reverse Primer | GGGACAGCGGTCATACGTG |
| ZFP423 | Forward Primer | TGGCCTGGGATTCCTCTGT |
| | Reverse Primer | CTCTTGACTTGTCACGCTGTT |
| ZFP467 | Forward Primer | TCCTGCTCAGGGCATGAGA |
| | Reverse Primer | TCCGAATCATCCATTCCTCCC |
| Tcf7l1 variant1 | Forward Primer | CGTCCCTGGTCAACGAATCG |
| | Reverse Primer | CTTCTCACTTCGGCGAAATAGT |
| Tcf7l1 variant2 | Forward Primer | ACGAGCTGATCCCCTTCCA |
| | Reverse Primer | CAGGGACGACTTGACCTCAT |
| KLF2 | Forward Primer | GAGCCTATCTTGCCGTCCTTT |
| | Reverse Primer | CACGTTGTTTAGGTCCTCATCC |
| Foxo1 | Forward Primer | CCCAGGCCGGAGTTTAACC |
| | Reverse Primer | GTTGCTCATAAAGTCGGTGCT |
| Foxa2 | Forward Primer | TCCGACTGGAGCAGCTACTAC |
| | Reverse Primer | GCGCCCACATAGGATGACA |
| Foxc2 | Forward Primer | AACCCAACAGCAAACTTTCCC |
| | Reverse Primer | GCGTAGCTCGATAGGGCAG |
| CD36 | Forward Primer | ATGGGCTGTGATCGGAACTG |

| | | |
|---|---|---|
| | Reverse Primer | TTTGCCACGTCATCTGGGTTT |
| Lipoprotein lipase | Forward Primer | ATGGATGGACGGTAACGGGAA |
| | Reverse Primer | CCCGATACAACCAGTCTACTACA |
| Fasn | Forward Primer | GGAGGTGGTGATAGCCGGTAT |
| | Reverse Primer | TGGGTAATCCATAGAGCCCAG |
| Plin1 | Forward Primer | CTGTGTGCAATGCCTATGAGA |
| | Reverse Primer | CTGGAGGGTATTGAAGAGCCG |
| Plin5 | Forward Primer | CTTCCTGCCCATGACTGAGG |
| | Reverse Primer | GACCCCAGACGCACAAAGTAG |
| Plin2 | Forward Primer | GACCTTGTGTCCTCCGCTTAT |
| | Reverse Primer | CAACCGCAATTTGTGGCTC |
| Plin3 | Forward Primer | ATGTCTAGCAATGGTACAGATGC |
| | Reverse Primer | CGTGGAACTGATAAGAGGCAGG |
| Plin4 | Forward Primer | GTGTCCACCAACTCACAGATG |
| | Reverse Primer | GGACCATTCCTTTTGCAGCAT |
| Plin5 variant1 | Forward Primer | TGTCCAGTGCTTACAACTCGG |
| | Reverse Primer | CAGGGCACAGGTAGTCACAC |
| Plin1 variant1 | Forward Primer | GGGACCTGTGAGTGCTTCC |
| | Reverse Primer | GTATTGAAGAGCCGGGATCTTTT |
| Dgat1 | Forward Primer | CTGATCCTGAGTAATGCAAGGTT |
| | Reverse Primer | TGGATGCAATAATCACGCATGG |
| Angptl4 | Forward Primer | CATCCTGGGACGAGATGAACT |
| | Reverse Primer | TGACAAGCGTTACCACAGGC |
| PDGFRα | Forward Primer | TATCCTCCCAAACGAGAATGAGA |

| | | |
|---|---|---|
| | Reverse Primer | GTGGTTGTAGTAGCAAGTGTACC |
| PDGFRβ variant1 | Forward Primer | AGGAGTGATACCAGCTTTAGTCC |
| | Reverse Primer | CCGAGCAGGTCAGAACAAAGG |
| PDGFRβ variant2 | Forward Primer | TTCCAGGAGTGATACCAGCTT |
| | Reverse Primer | AGGGGGCGTGATGACTAGG |
| VEGFc | Forward Primer | GTGAGGTGTGTATAGATGTGGGG |
| | Reverse Primer | ACGTCTTGCTGAGGTAACCTG |
| VEGFb | Forward Primer | GCCAGACAGGGTTGCCATAC |
| | Reverse Primer | GGAGTGGGATGGATGATGTCAG |
| EGR2 | Forward Primer | CCAGACACTACTTTCCACCG |
| | Reverse Primer | CCTCCCAGTTCACCATTGGG |
| Resistin | Forward Primer | CAGGTCGCTTCCTGATGTCG |
| | Reverse Primer | CCAGACCCTCAGCTTAGACC |
| Lpl(154) | Forward Primer | AGGCATACAGGTGCAACTCC |
| | Reverse Primer | TAGGGCATCTGAGAGCGAGT |
| Fabp4 | Forward Primer | TAAAAGTGAGCTATCTGGACTTCA |
| | Reverse Primer | GCCCACTCCCACTTCTTTCAT |
| CREB1 | Forward Primer | GAGAAGCGGAGTGTTGGTGA |
| | Reverse Primer | ACTCTGCTGGTTGTCTGCTC |
| TET1 variant1 | Forward Primer | GCAGTGAACCCCGGAAAAC |
| | Reverse Primer | AGAGCCATTGTAAACCCGTTG |
| TET1 variant2 | Forward Primer | ACACAGTGGTGCTAATGCAG |
| | Reverse Primer | AGCATGAACGGGAGAATCGG |
| TET2 | Forward Primer | CTCCCATCAGCCATACAGAACC |

| | | |
|---|---|---|
| | Reverse Primer | CTGACTGTGCGTTTTATCCCT |
| TET3 | Forward Primer | TGCGATTGTGTCGAACAAATAGT |
| | Reverse Primer | TCCATACCGATCCTCCATGAG |
| ADIPOQ | Forward Primer | ATCTGGAGGTGGGAGACCAA |
| | Reverse Primer | GGGCTATGGGTAGTTGCAGT |
| GATA2 | Forward Primer | CACCCCGCCGTATTGAATG |
| | Reverse Primer | CCTGCGAGTCGAGATGGTTG |
| EBF1 | Forward Primer | AAGCATCCAACGGAGTGGAAG |
| | Reverse Primer | GATTTCCGCAGGTTAGAAGGC |
| Hoxa6 | Forward Primer | CGGCCAGGACTCCTTCTTG |
| | Reverse Primer | CCGAGTTGGACTGTTGGTAAAA |
| Hoxa5 | Forward Primer | CTCATTTTGCGGTCGCTATCC |
| | Reverse Primer | ATCCATGCCATTGTAGCCGTA |
| Vdr | Forward Primer | GAATGTGCCTCGGATCTGTGG |
| | Reverse Primer | ATGCGGCAATCTCCATTGAAG |
| KLF4 | Forward Primer | GGCGAGTCTGACATGGCTG |
| | Reverse Primer | GCTGGACGCAGTGTCTTCTC |
| ATF7 | Forward Primer | ATCATTGCAGATCAAACGCCT |
| | Reverse Primer | AGCACCCTTTTTCTCATCGTC |
| JunB | Forward Primer | TCACGACGACTCTTACGCAG |
| | Reverse Primer | CCTTGAGACCCCGATAGGGA |
| Pbx1 Variant a | Forward Primer | CAGCGGGTTCTTCCAGTTCTT |
| | Reverse Primer | CGAGTCCGTCACTGTATCCTC |
| Pbx1 Variant b | Forward Primer | TGAAGCCTGCCTTGTTTAATGT |

| | | |
|---|---|---|
| | Reverse Primer | ATGTTGTCCAGTCGCATGAGC |
| Slc2a1 | Forward Primer | TACACCCCAGAACCAATGGC |
| | Reverse Primer | CCCGTAGCTCAGATCGTCAC |
| Lyz2(NM_017372.3) | Forward Primer | AGCACACTGTCAAAACCGAG |
| | Reverse Primer | CTTAGAGGGGAAATCGAGGGAA |
| Lyz2(17105) | Forward Primer | TGCCAGAACTCTGAAAAGGAATGG |
| | Reverse Primer | CAGTGCTTTGGTCTCCACGGTT |
| GM14325 (NM_001024849.3) | Forward Primer | CCATAAATGCATTCAATATGGGGA |
| | Reverse Primer | ACATTTCTTGTATCCCCTTGAACT |
| GM10354 (NM_001281514.1) | Forward Primer | CAAAGAGATCCAGCTCACTATGGA |
| | Reverse Primer | GGGATTTTGCCTGTGGTAGGG |

**A)**

| Samples | Raw reads | Filter and Trim reads | Clean ratio | Mapped reads | Mapping Rate (%) | Duplicated reads | Duplicated Rate (%) |
|---|---|---|---|---|---|---|---|
| Adipocyte | 870974927 | 870967170 | 99.07 | 648845222 | 74.5 | 63739281 | 9.82 |
| Preadipocyte | 891784966 | 891779716 | 99.07 | 666335293 | 74.72 | 61245031 | 9.19 |

**B)**

| Samples | Total Cytosine | Methylated CpG | Unmethylated CpG | mCpG(%) | mCHG(%) | mCHH(%) |
|---|---|---|---|---|---|---|
| Adipocyte | 24688537793 | 797818959 | 454925272 | 63.69 | 0.13 | 0.18 |
| Preadipocytes | 25172225618 | 834519959 | 449432824 | 65 | 0.11 | 0.15 |

**Table S3**: Summary of AR WGBS dataset and read counts in pre-adipocytes and adipocytes depicting increased methylated CpGs as the pre-adipocytes differentiated. A) Mapping of raw reads with reference genome mm10. B) Total analysis of methylated cytosine showing CG cytosines and non-CG cytosines.

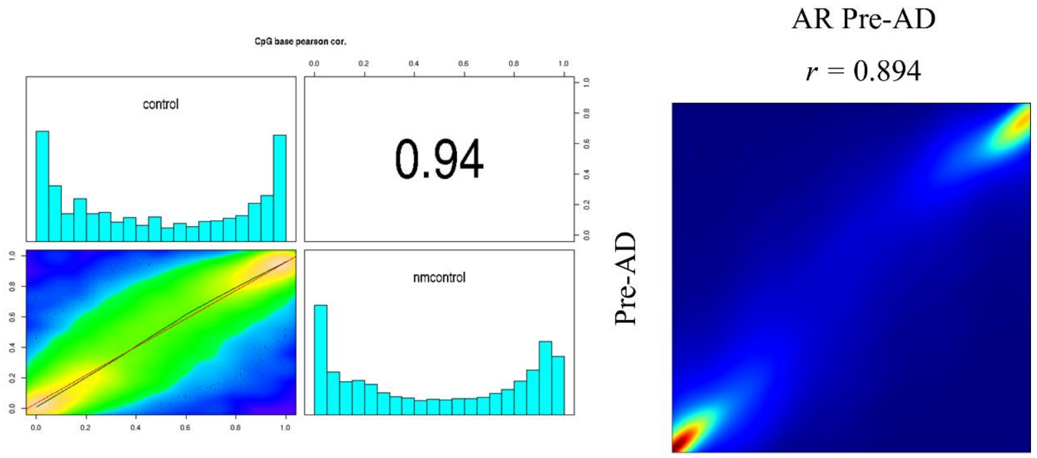


**Figure S6**: Pearson Correlation between NABI control vs. AR control. Total locations of NABI control with coverage 10:29584. Total locations of AR pre-adipocyte with coverage 10: 32860286. Common locations used for correlation analysis: 28973.

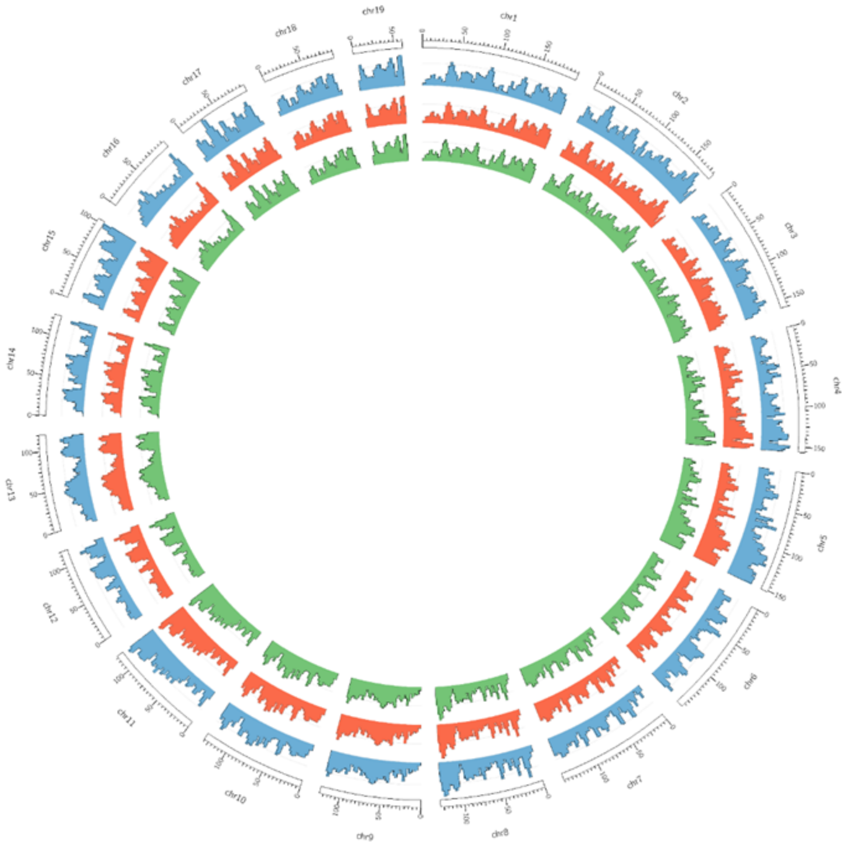


**Figure S7**: Comparison of NABI & AR data. Red color indicates pre-adipocytes and green represents the adipocytes whereas blue indicates NABI control.

| **Read Depth** | **Correlation(NABI vs AR)** | **Common CpG locations** |
|---|---|---|
| 4 | 0.866957 | 2830772 |
| 5 | 0.8808512 | 1057385 |
| 6 | 0.89205 | 379097 |
| 7 | 0.9050764 | 141414 |
| 8 | 0.9194332 | 63271 |
| 9 | 0.9312149 | 37840 |
| 10 | 0.9397303 | 28973 |
| 11 | 0.9437549 | 25454 |
| 12 | 0.9469991 | 23563 |

**Table S4**: Correlation analysis at various depths (from 4 to 12) for NABI control vs AR control.

| Hyper 15D | Hyper 2D | Hypo 15D | Hypo 2D |
|---|---|---|---|
| Negative regulation of cytokine production | mucopolysaccharide metabolic process | erythrocyte differentiation | ruffle assembly |
| nucleoside monophosphate biosynthetic process | mitochondrial transmembrane transport | regulation of fat cell differentiation | apoptotic process involved in the development |
| positive regulation of interleukin-12 production | nitric oxide biosynthetic process | regulation of protein localization to the nucleus | regulation of extrinsic apoptotic signaling pathway in the absence of ligand |
| cellular detoxification | regulation of osteoclast differentiation | establishment of cell polarity | spliceosomal complex assembly |
| glycogen biosynthetic process | cellular response to unfolded protein | regeneration | response to light stimulus |
| glucan biosynthetic process | regulation of multicellular organism growth | kidney epithelium development | response to pH |
| positive regulation of mitotic nuclear division | proteoglycan metabolic process | regulation of osteoblast differentiation | apoptotic cell clearance |
| intestinal absorption | nitric oxide metabolic process | nephron development | positive regulation of G1/S transition of mitotic cell cycle |
| cellular response to lipopolysaccharide | negative regulation of myeloid cell differentiation | erythrocyte homeostasis | negative regulation of the production of molecular mediators of immune response |
| regulation of macrophage cytokine production | intermediate filament cytoskeleton organization | negative regulation of the Wnt signaling pathway | mitotic spindle checkpoint signaling |
| positive regulation of protein tyrosine kinase activity | killing of cells of another organism | regulation of ubiquitin-dependent protein catabolic process | establishment of protein localization to the endoplasmic reticulum |
| regulation of polysaccharide metabolic process | intermediate filament-based process | cellular response to starvation | negative regulation of intracellular protein transport |

| | | | |
|---|---|---|---|
| toll-like receptor 4 signaling pathway | sulfur compound biosynthetic process | signal transduction in response to DNA damage | regulation of TORC1 signaling |
| positive regulation of signaling receptor activity | regulation of oxidoreductase activity | BMP signaling pathway | regulation of fibroblast migration |
| cellular response to molecule of bacterial origin | establishment of protein localization to mitochondrion | protein import into nucleus | regulation of peptidyl-threonine phosphorylation |
| mRNA polyadenylation | cyclic-nucleotide-mediated signaling | regulation of response to wounding | biological phase |
| positive regulation of the glucose metabolic process | inclusion body | hindbrain development | peptidyl-serine phosphorylation |
| apoptotic cell clearance | Golgi stack | ruffle | negative regulation of dephosphorylation |
| negative regulation of I-kappa B kinase/NF-kappaB signaling | cortical actin cytoskeleton | focal adhesion | regulation of alcohol biosynthetic process |
| peptidyl-tyrosine phosphorylation | antioxidant activity | carboxylic acid binding | spindle checkpoint signaling |
| RNA polyadenylation | | phosphatidylinositol phosphate binding | regulation of hormone metabolic process |
| peptidyl-tyrosine modification | | | DNA geometric change |
| regulation of transcription from RNA polymerase II promoter in response to stress | | | regulation of erythrocyte differentiation |
| positive regulation of nitric oxide biosynthetic process | | | regulation of intracellular transport |
| cellular response to toxic substance | | | phospholipid translocation |
| regulation of ERK1 and ERK2 cascade | | | regulation of mitotic cell cycle phase transition |

| | | | |
|---|---|---|---|
| Golgi apparatus subcompartment | | | regulation of humoral immune response |
| scavenger receptor activity | | | positive regulation of glucose transmembrane transport |
| MHC class I protein binding | | | regulation of ubiquitin-protein transferase activity |
| intramembrane lipid transporter activity | | | negative regulation of protein phosphorylation |
| SH2 domain binding | | | lipid translocation |
| general transcription initiation factor activity | | | positive regulation of glial cell differentiation |
| | | | histone H3-K9 modification |
| | | | neurotransmitter receptor complex |
| | | | chloride channel complex |
| | | | integral component of mitochondrial inner membrane |
| | | | intrinsic component of mitochondrial inner membrane |
| | | | cyclin-dependent protein kinase holoenzyme complex |
| | | | disulfide oxidoreductase activity |

**Table S5:** Comparison of methylation-dependent gene expression analysis of DMRs present at 2D and 15D; 15D depicting increased expression of certain pathways involved in dedifferentiation.

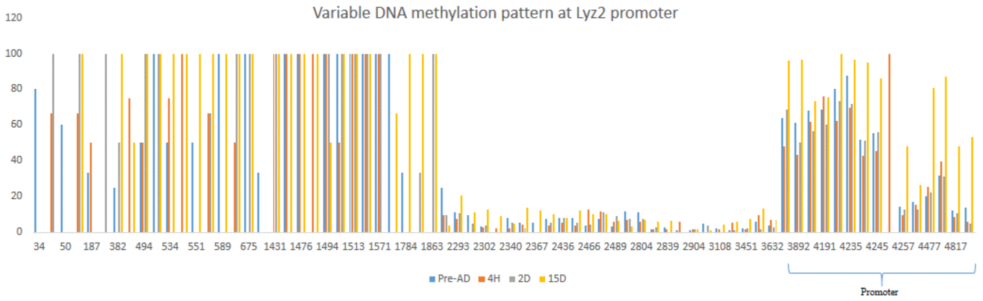


**Figure S9:** Hypermethylation at Lyz2 promoter at terminally differentiated adipocytes.

**Supplementary References:**


1. J. Park *et al.*, "Targeted erasure of DNA methylation by TET3 drives adipogenic reprogramming and differentiation," *Nature Metabolism*, vol. 4, no. 7, pp. 918–931, 2022.
2. Emmert Buck *et al.*, "Quantitative RT-PCR gene expression analysis of laser micro-dissected tissue samples," *Nature Protocols,* vol. 4, pp. 902–922, 2009.
